# A multifaceted architectural framework of the mouse claustrum complex

**DOI:** 10.1101/2022.06.02.494429

**Authors:** Joachim S. Grimstvedt, Andrew M. Shelton, Anna Hoerder-Suabedissen, David K. Oliver, Christin H. Berndtsson, Stefan Blankvoort, Rajeevkumar R. Nair, Adam M. Packer, Menno P. Witter, Clifford G. Kentros

## Abstract

Accurate anatomical characterizations are necessary to investigate neural circuitry on a fine scale, but for the rodent claustrum complex (CC) this has yet to be fully accomplished. The CC is generally considered to comprise two major subdivisions, the claustrum (CL) and the dorsal endopiriform nucleus (DEn), but regional boundaries to these areas are highly debated. To address this, we conducted a multifaceted analysis of fiber- and cyto-architecture, genetic marker expression, and connectivity using mice of both sexes, to create a comprehensive guide for identifying and delineating borders to the CC. We identified four distinct subregions within the CC, subdividing both the CL and the DEn into two. Additionally, we conducted brain-wide tracing of inputs to the entire CC using a transgenic mouse line. Immunohistochemical staining against myelin basic protein (MBP), parvalbumin (PV), and calbindin (CB) revealed intricate fiber-architectural patterns enabling precise delineations of the CC and its subregions. Myelinated fibers were abundant in dorsal parts of the CL but absent in ventral parts, while parvalbumin labelled fibers occupied the entire CL. Calbindin staining revealed a central gap within the CL, which was also visible at levels anterior to the striatum. Furthermore, cells in the CL projecting to the retrosplenial-cortex were located within the myelin sparse area. By combining our own experimental data with digitally available datasets of gene expression and input connectivity, we could demonstrate that the proposed delineation scheme allows anchoring of datasets from different origins to a common reference framework.

**Significance statement:** Mice are a highly tractable model for studying the claustrum complex (CC). However, without a consensus on how to delineate the CC in rodents, comparing results between studies is challenging. It is therefore important to expand our anatomical knowledge of the CC, to match the level of detail needed to study its functional properties. Using multiple strategies for identifying claustral borders, we created a comprehensive guide to delineate the CC and its subregions. This anatomical framework will allow researchers to anchor future experimental data into a common reference space. We demonstrated the power of this new structural framework by combining our own experimental data with digitally available data on gene expression and input connectivity of the CC.

## Introduction

To conduct fine-scale functional investigations of brain circuitry, we require accurate anatomical frameworks to integrate observations across studies. Perhaps no brain area is more in need of this than the rodent claustrum complex (CC), where regional borders have been debated for decades. There is a rapidly growing interest in understanding the functional roles of the CC, as it may be involved in key elements of cognition like attention (Atlan et al., 2018; Goll et al., 2015), multimodal integration (Crick & Koch, 2005; Shelton et al., 2022), and memory processing (Behan & Haberly, 1999; Witter et al., 1988), processes that likely depend on the extensive connectivity between the CC and prefrontal, sensory, and parahippocampal regions (Atlan et al., 2017; Zingg et al., 2018). Generally, the CC is considered to comprise two main subdivisions, the claustrum (CL) and the dorsal endopiriform nucleus (DEn) (Kowianski et al., 1999), however, it is challenging to distinguish these two regions from one another and the adjacent cortex. To remedy this, we studied the combinatorial expression patterns of multiple markers to develop a multifaceted anatomical description of CC.

Patterns in fiber-architecture can be used to identify regional boundaries (Glasser & Van Essen, 2011). For example, the primate CC can be clearly distinguished from surrounding cortices due to the extreme and external capsules (Berman et al., 2020; Pham et al., 2019). In rodents the extreme capsule is only rudimentary, making the CC, and especially the CL, difficult to distinguish from cortex (Bruguier et al., 2020; Kowianski et al., 1999). However, myelinated axons do surround CL in mice, whereas CL itself lacks myelination (Wang et al., 2017; Wang et al., 2022). A well-established landmark for CL is the plexus formed by parvalbumin (PV) positive neurites (Druga et al., 1993; Real et al., 2003). Conversely, calbindin-D28k (CB) shows reduced labeling in CL (Celio, 1990). Together, these markers may provide a good fiber and neurochemical foundation to distinguish the CL from the rest of the CC.

Brain regions can also be defined by connectivity patterns. An example of this is the pocket of retrosplenial-cortex (RSC) projecting cells located centrally in CL, roughly in the middle of the PV-plexus (Marriott et al., 2021; Zingg et al., 2018). This organization is often referred to as the ‘core and shell’ model of CL (Atlan et al., 2017; Davila et al., 2005; Marriott et al., 2021; Real et al., 2003), although the existence of a CL shell has been debated (Mathur, 2014). Additionally, dorsoventral gradients in claustral input connectivity have been reported (Atlan et al., 2017; Olson & Graybiel, 1980; Witter et al., 1988). As such, it may be that connectivity by itself is not sufficient to define CC.

The functional and ontogenetic relationship between the CL and DEn in rodents has long been a matter of debate (Binks et al., 2019; Watson & Puelles, 2017). We follow the recent proposal to classify these regions as subdivisions of the same complex, based on anatomical and genetic similarities (Smith et al., 2019). Several genetic markers show elevated expression in both the CL and DEn (Watakabe et al., 2014), while others show subregional specificity (Bruguier et al., 2020; Dillingham et al., 2017; Dillingham et al., 2019; Erwin et al., 2021; Watson & Puelles, 2017). In rodents, the DEn has not been studied as extensively as the CL but exhibits distinct patterns in connectivity and neuroanatomical markers (Beneyto & Prieto, 2001; Suzuki & Bekkers, 2010; Witter et al., 1988).

We present a comprehensive guide for delineating the mouse CC based on patterns in fiber- and cyto-architecture. Next, we relate this framework with a combination of our own experimental data on brain-wide inputs to CL and DEn and publicly available gene expression and input connectivity data. Together, these patterns revealed what we consider a new, robust, and versatile definition of the mouse CC and its subdivisions that will be of use to future functional studies on this region.

## Materials and methods

### Animal care and husbandry

Experiments were conducted at the Kavli Institute for Systems Neuroscience at the Norwegian University of Science and Technology (NTNU), Trondheim, and the Department of Physiology, Anatomy & Genetics, at the University of Oxford. Animals were group housed in environmentally enriched cages and given *ad libitum* access to food and drink. Animals housed at the University of Oxford were kept at a normal 12:12h day/night cycle, while at NTNU the day/ night cycle was reversed. Both male and female animals were used. All procedures involving animals were done in accordance with guidelines of the Federation of European Laboratory Animal Science Association (FELASA), and local authorities at NTNU and the University of Oxford. Surgical procedures performed at the University of Oxford were carried out under license from the UK Home Office in accordance with the Animal (Scientific Procedures) Act 1986. Experiments carried out at NTNU were approved by the Norwegian Food Safety Authority (FOTS).

### Transgenic mouse lines

Two transgenic mouse lines were used in this study: the claustrum complex – enhancer driven gene expression (CC-EDGE) line and Tre-Tight-THAG line. The cross between these two, *CC-EDGE::TRE-Tight-THAG*, was used in monosynaptic rabies tracing experiments. The CC-EDGE transgenic mouse line, originally called MEC-13-53D (Blankvoort et al., 2018), expresses tetracycline transactivator (tTA) protein within a subpopulation of cells that are largely confined to the CC. The Tre-Tight-THAG transgenic line was generated using the following steps. The gene sequences of avian-specific tumor virus receptor A (TVA), hemagglutinin tag (HA) and challenge virus standard-11 glycoprotein (CVS11G) separated by 2A elements were inserted between Xma1 and Mlu1 restriction sites in the pTT2 construct previously described in Weible A.P et al (2010). After the sequence verification, the resulting construct pTT2-TVA-2A-2xHA-2A-CVS11G was linearized, run on 1% agarose gel, and purified using Zymoclean Gel DNA Recovery Kit (Zymo research, D4001) as per protocol. The transgenic mouse facility of the University of Oregon carried out pronuclear injections to create the TRE-Tight-THAG line. By crossing the CC-EDGE line to the TRE-Tight-THAG transgenic mouse line, rabies glycoprotein and TVA receptors were conditionally expressed in tTA producing cells.

### General surgical procedures and tissue acquisition

General anesthesia was induced with 5% isoflurane (IsoFlo® vet) prior to surgical procedures. A continuous flow of isoflurane and oxygen was administered throughout surgeries and adjusted to maintain stable anesthesia. Animals were placed in a stereotaxic frame while resting on a heating pad at 37°C for the duration of the procedure. Analgesics were given prior to surgery, either through subcutaneous injections of metacam (1 mg/kg) and temgesic (0.1 mg/kg) or intraperitoneal injections of metacam (5 mg/kg) and buprenorphine (0.1 mg/kg). Saline injections were administered during the surgery to avoid dehydration. The incision area was disinfected with iodine and subcutaneously injected with a local anesthetic (Marcaine® 1 mg/kg or bupivacaine) for 2 minutes before the initial incision. The cranium was manually leveled between bregma and lambda, and between the left and right hemisphere. Craniotomies were performed using a dental drill and were exclusively done in the right hemisphere unless otherwise noted. Pulled glass pipettes were used for all injections. Dorsoventral depth was measured from the surface of the pia. Pipettes remained in place for 10 minutes before retraction. Postoperative analgesia was administered 7-12h after surgery, and animals were given easily ingestible food (oat porridge). Further analgesia was administered when necessary. Prior to tissue collection, animals were deeply anaesthetized with isoflurane, and given a lethal intraperitoneal injection of pentobarbital (0.1ml ip of a solution of 100mg/ml). Animals were carefully observed until breathing ceased, and motor and eye blinking reflexes were gone, at which point transcardial perfusion was performed using a peristaltic pump (MasterFlex®, USA) to pump Ringer saline solution followed by 4% paraformaldehyde (Sigma Aldrich) in 0.125M phosphate buffer through the circulatory system. Brains were post-fixed in 4% paraformaldehyde overnight and then transferred to a cryo-protective solution containing 20% glycerol and 2% dimethyl sulfoxide (DMSO) in 0.125M phosphate buffer.

### Histology and immunohistochemistry

Brains were sliced into 40µm thick coronal sections using a freezing sliding-microtome (Thermo Scientific™ HM 430, USA), kept at approximately -40°C. Four equally spaced series of sections were collected for each brain, allowing for multiple histological staining procedures with the same brain. Various immunohistochemical (IHC) procedures were carried out with standard protocols for free floating brain sections. Permeabilization of brain sections was done using a phosphate buffer solution containing 0.5% TritonX-100 (Merck KGaA). Blocking was done with 5% normal goat serum (Abcam, #ab7481) at room temperature. Primary antibody incubation was done for at least 48 hours at 4-5 °C. Secondary antibody incubation was done for 1-2 hours at room temperature. Sections were mounted onto microscope slides (Menzel-Gläser SuperFrost®Plus) and left to dry overnight. On the following day, sections were cleared for 10 minutes in Toluene and coverslipped in a mixture of Toluene and Entellan (Merck KGaA). Another mounted series was used for Nissl staining with Cresyl Violet (Sigma® Life Science, C5042), after tissue clearing in Xylene (Merck KGaA) and multiple steps of rehydration from 100-50% ethanol. Some experiments involved IHC staining followed by de-coverslipping and Nissl staining with Cresyl Violet. We tested several markers to label myelinated axons and settled on using an antibody against myelin basic protein (MBP; Merck, NE1019-100UL, 1:1000). We used two different antibodies for both PV (Sigma, P3088, 1:1000; Swant, PV27, 1:1000) and CB (Swant, 300, 1:3000; Swant, CB38, 1:3000), depending on host-organism availability in each procedure. The extent of neurite labeling in PV and CB staining varied considerably with the choice of primary and secondary antibody. Two animals stained against MBP and CB were originally used to characterize the crossbreed of transgenic lines CC-EDGE and tetO-Chrimson, but as they showed no apparent variation in expression of these markers they were included in this dataset. Alexa Fluor-conjugated secondary antibodies (Thermo Fischer Scientific, Invitrogen, 1:400) were used in all IHC procedures.

### Neuroanatomical tract tracing

Two different injection strategies were used in this study (Table 1). *CC-EDGE::TRE-Tight-THAG* transgenic mice (aged 11-23 weeks) were unilaterally injected with EnvA-pseudotyped, G-protein deleted CVS-N2c recombinant rabies virus expressing tdTomato (RABV-tdTomato, 107 functional virus particles/ml) at one or two coordinates in the claustrum complex. The RABV-tdTomato virus was provided by Dr. Rajeevkumar R. Nair and produced at the Viral Vector Core at the Kavli Institute for Systems Neuroscience. RABV-tdTomato was injected at 10-20μl/ sec. The brains from these animals were collected 10-14 days after the injection following the procedure described above. C57BL6J mice (aged 3-5 weeks) were unilaterally injected with Cholera Toxin Subunit B (CTB (Recombinant) Alexa Fluor™ 647 Conjugate [0.1% wt/vol, Thermo Fisher C34778]) in the retrosplenial cortex. CTB was injected at 5-10nl/sec. CTB injected animals were allowed to recover for a minimum of 5 days before brains were collected.

**Table 1:** Injection coordinates

| Animal ID | Injection volume (nl) | Sex | Injection coordinates (mm) |  |  |
| --- | --- | --- | --- | --- | --- |
|  |  |  | Anteroposterior | Mediolateral | Dorsoventral |
| Rabies tracing |  |  |  |  |  |
| Rb1 | 750/750 | F | +1.33/+0.11 | +3.2/+4.0 | -2.8/-2.8 |
| Rb2 | 500/600 | F | +1.33/+0.11 | +3.4/+4.1 | -2.9/-2.8 |
| Rb3 | 500/500 | F | +1.33/+0.11 | +3.2/+4.0 | -2.9/-2.8 |
| Rb4 | 500/500 | M | +1.33/+0.11 | +3.2/+4.0 | -2.9/-2.8 |
| Rb5 | 150/150 | F | +1.33/+0.11 | +3.2/+4.1 | -2.9/-3.1 |
| Rb6 | 800 | M | +1.33 | +3.2 | -2.9 |
| Rb7 | 600/600 | F | +1.33/+0.11 | +3.2/+4.0 | -2.8/-2.8 |
| Rb8 | 500/500 | M | +1.33/+0.11 | +3.2/+4.3 | -2.9/-2.8 |
| Rb9 | 300/300 | F | +1.33/+0.11 | +3.1/+3.7 | -3.2/-3.2 |
| Rb10 | 400/200 | M | +1.33/+0.11 | +3.1/+3.8 | -3.2/-3.2 |
| Rb11 | 800 | F | +1.33 | +3.1/+3.7 | -3.2/-3.2 |
| Rb12 | 200 | M | +1.33 | +3.3 | -2.6/-2.6 |
| Rb13 | 100 | F | -0.7 | -0.7* | -2.6/-2.6 |
| CTB tracing |  |  |  |  |  |
| CTB1 | 80 | M | -3.0 | +0.5 | -1.0 |
| CTB2 | 80 | M | -3.0 | +0.5 | -1.0 |
| CTB3 | 80 | M | -3.0 | +0.5 | -1.0 |
| CTB4 | 80 | M | -3.0 | +0.5 | -1.0 |
| CTB5 | 80 | M | -3.0 | +0.5 | -1.0 |
| CTB6 | 80 | M | -3.0 | +0.5 | -1.0 |

\*Measured from the lateral edge of the cranium

### Quantification of brain-wide monosynaptic inputs to the claustrum complex

Rabies injected brains were immunohistochemically stained using antibodies against the 2A linker protein (Merck/ Millipore, ABS31, 1:2000) found in tTA expressing cells, and a red fluorescent protein tag (tdTomato; ChromoTek, 5f8-100, 1:500) expressed by the rabies virus. Cell counting was done with Neurolucida 2014 software (MBF Bioscience). Following cell quantification, each slide was decoverslipped and stained with Cresyl Violet. Images of each section were then superimposed onto the contours of the sections and the position markers of counted cells, allowing close approximation of the fluorescent and cytoarchitectural images resulting in the accurate position of input cells throughout the brain. Microsoft Excel and custom-made scripts in Matlab were used to analyze and visualize the quantification of the tracing data.

### Image acquisition and processing

Fluorescence and brightfield images were acquired using a slide scanner with a Plan-Apochromat 20X/0.8 NA M27 objective, resulting in a resolution of 0.325 µm/pixel (Axio Scan.Z1, ZEISS). A series of higher resolution fluorescent images were obtained with a confocal microscope (LSM 880, ZEISS). Images were post-processed with ZEN Black 2.1 SP2, ZEN Blue 2.3 Lite, and Adobe Photoshop to enhance the signal quality. All image processing was applied to the entire image. CTB images were acquired using an Olympus FV3000 confocal laser scanning microscope and post-processed in ImageJ and Python 3.7.

### Rostrocaudal landmarks

We selected 5 landmarks to locate different rostrocaudal levels of the CC. The landmarks were approximately at the same dorsoventral level as the CC, to mitigate variation from slight differences in the sectioning plane. Distance to bregma was estimated by comparison to a reference atlas (Paxinos & Franklin, 2019). The far rostral landmark was selected where the orbital cortex merges with the olfactory cortex (B+2.09). As the rostral landmark, we selected the caudal-most section displaying the dorsal peduncular area, positioned between the corpus callosum and the septal complex (B+0.97). Roughly at the midpoint of the CC, the central landmark was chosen as the point where the anterior commissure joins at the midline (B+ 0.13). The caudal landmark was selected at the rostral-most section showing the basolateral amygdala (B-0.59). Finally, the far caudal landmark was chosen where the optic tract joins with the internal capsule (B-1.43). Notably, the far rostral and far caudal landmarks do not represent the rostral and caudal edges of the CC.

### Delineation references

We used a combination of research articles and atlases for our cyto-architectural delineations of brain regions. Table 2 lists articles that were used, and for which borders they were used. Other regions were delineated based on the Paxinos & Franklin Mouse reference atlas (Paxinos & Franklin, 2019).

**Table 2:**
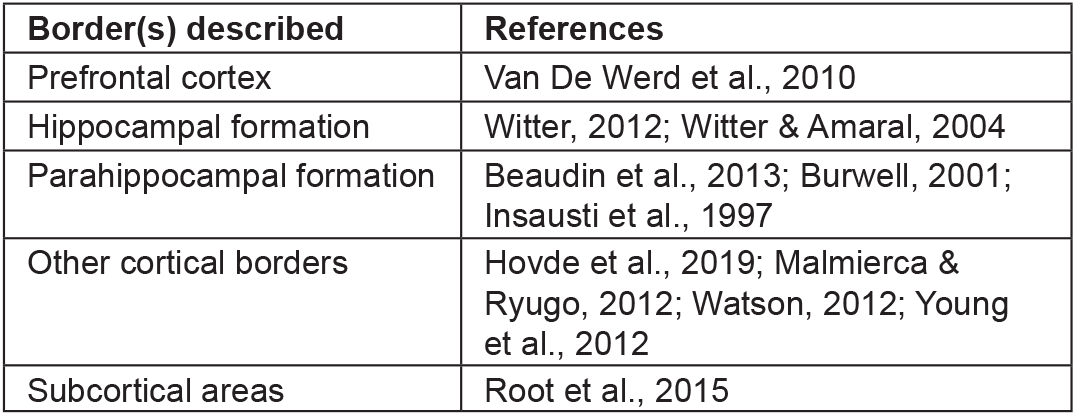
Delineation references

| Border(s) described | References |
| --- | --- |
| Prefrontal cortex | Van De Werd et al., 2010 |
| Hippocampal formation | Witter, 2012; Witter & Amaral, 2004 |
| Parahippocampal formation | Beaudin et al., 2013; Burwell, 2001; Insausti et al., 1997 |
| Other cortical borders | Hovde et al., 2019; Malmierca & Ryugo, 2012; Watson, 2012; Young et al., 2012 |
| Subcortical areas | Root et al., 2015 |

### Data-approximation by image-warping

Images were collected from online *in situ* hybridization (ISH) and tracer databases (© 2006 Allen Institute for Brain Science, ISH Data, Available from: https://mouse.brain-map.org/, © 2017 Allen Institute for Brain Science, Projection Dataset, Available from: https://connectivity.brain-map.org/), and used in an image approximation procedure. References to individual experiments are given in tables 3 and 7. The BigWarp function in ImageJ was used to align each brain section to a reference by creating a set of transformation coordinates (Bogovic et al., 2016; Schneider et al., 2012). An alignment frame was drawn onto each section using vector graphics in Adobe Illustrator, to assign the necessary number of anchor points to allow for accurate transformation. A few cortical borders were indicated, and equidistant points between these borders were then marked to divide the cortex into multiple “bins” (n=4 bins for insular cortex, n=4 bins for piriform cortex and n=8 bins from somatosensory cortex to cingulate cortex). ISH data were thresholded using the inbuilt function in ImageJ, to remove background and allow the data to be superimposed onto the reference image.

**Table 3:**
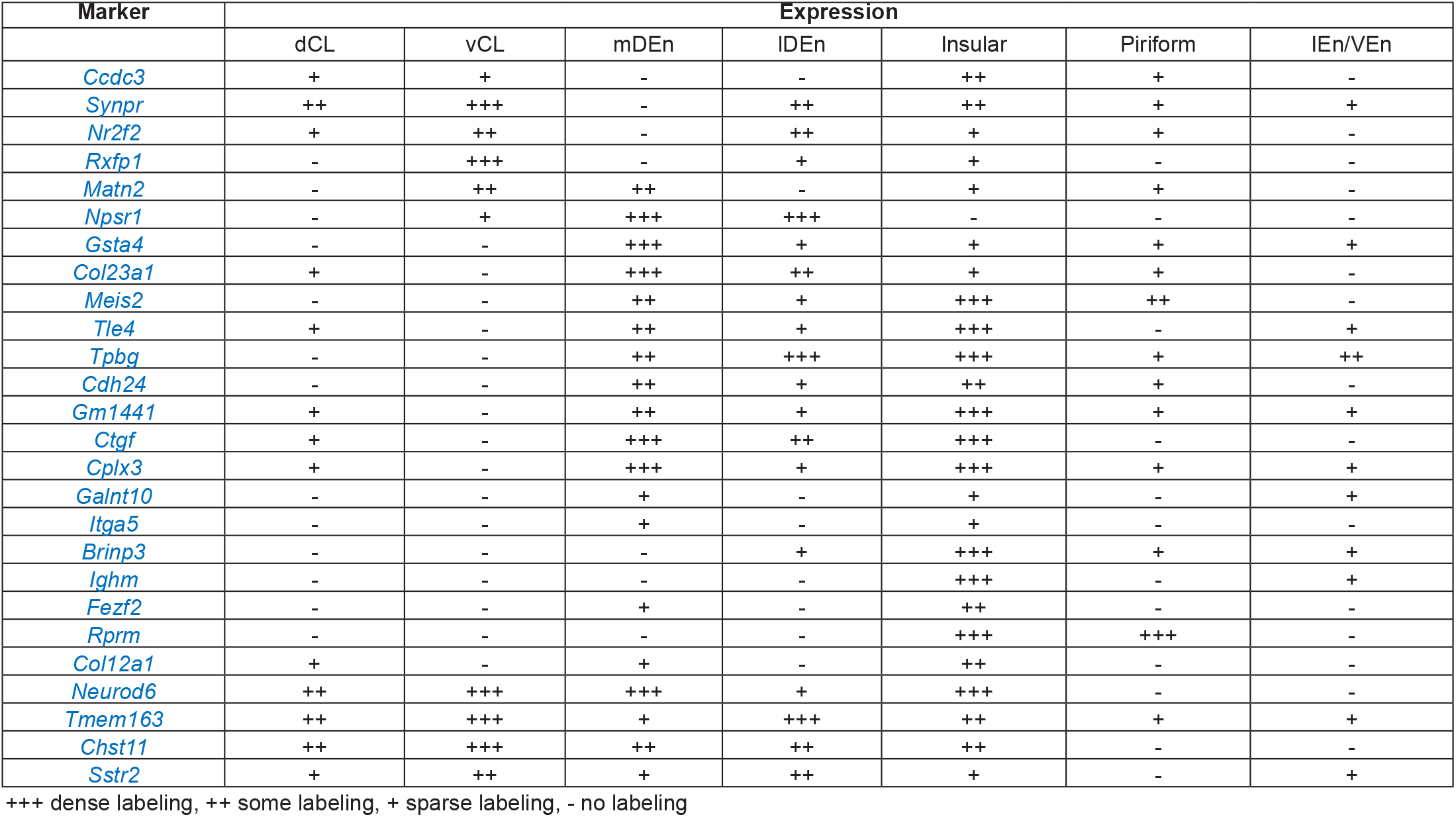
Genetic marker candidates for the claustrum complex

### Fluorescence profile analysis

Average fluorescence traces were measured in ImageJ using the “Plot profile” function to produce average signal intensity traces within a rectangular region. In fiber-architectural comparisons (Figure 1F), bilateral claustra were analyzed in each animal (n=5 animals stained against PV and MBP, n=3 animals stained against CB). For these measurements, a rectangular area was placed in approximately the same location in each section, as determined by the image warping pipeline described above. For Figure 5, images across mice (n=6) and within matched representative sections were aligned to the max CTB signal, cropped to the same area and rotated along the external capsule. Images were then normalized and averaged along the horizontal and vertical axes to produce fluorescence traces. The fluorescence profiles were processed in Matlab or Python and smoothed using a Gaussian filter. The intensity of the signal was scaled using min-max normalization.

**Figure 1:**
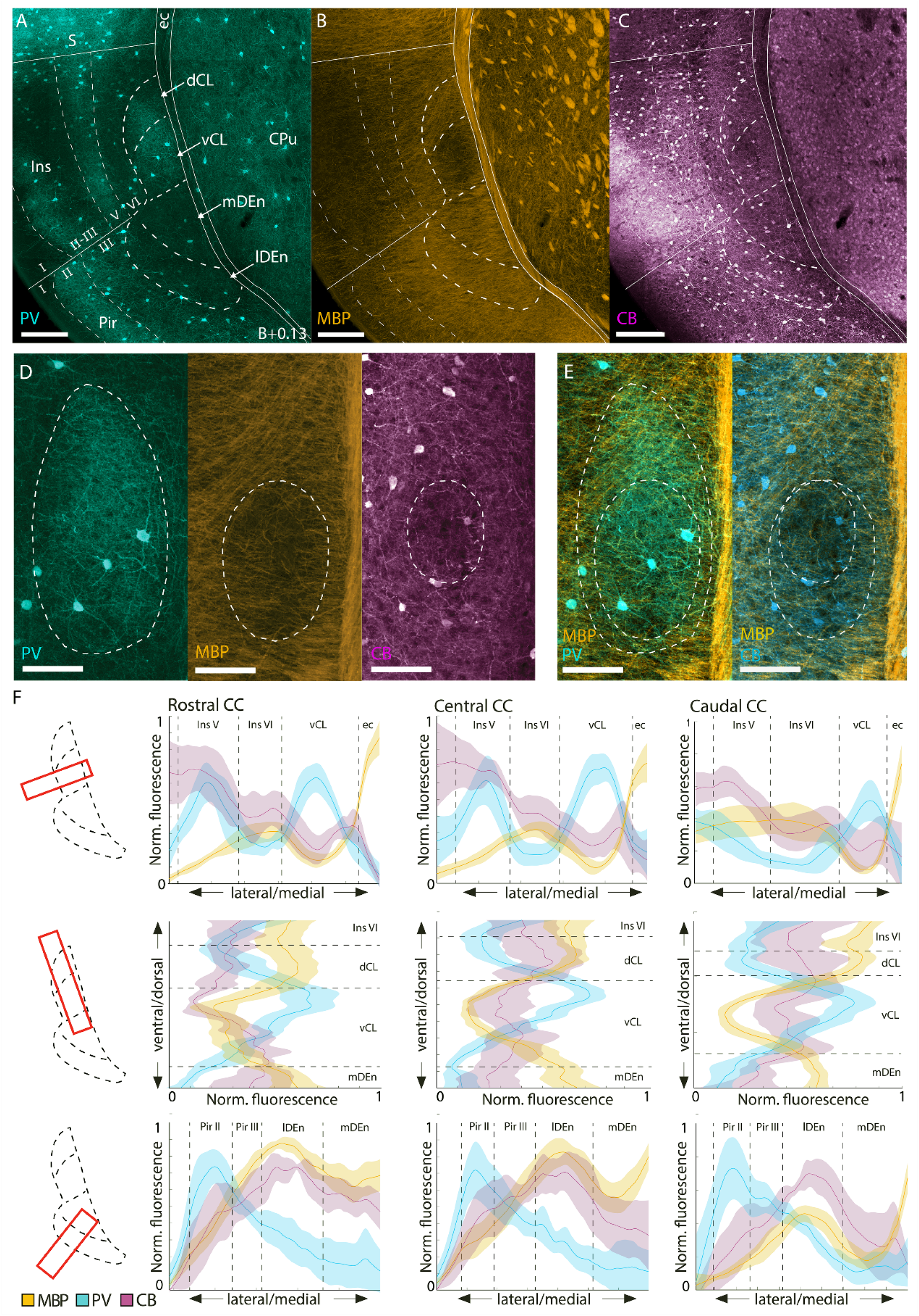
Fiber-architectural characterization of the claustrum complex. **A-C)** Reappraisal of borders to CC based on overlapping patterns in tissue triple-stained for parvalbumin (PV) myelin-basic-protein (MBP), and calbindin (CB). Scale bars measure 200μm. **D)** Delineations of the PV-plexus, MBP-gap, and CB-gap in the same tissue. **E)** Merged image panels from D showing MBP/PV and MBP/CB expression. Both PV and CB labeling are pseudo-colored cyan to increase visibility. Scale bars measure 100μm. **F)** Average fluorescence traces of PV (n=5 mice) MBP (n=5 mice), and CB (n=3 mice) along mediolateral and dorsoventral gradients, at rostral (B+0.97), central (B+0.13) and caudal (B-0.59) levels of the CC. Bilateral measurements were made in each brain. Dashed lines indicate approximate borders based on the combined profile patterns. Fluorescence measures were normalized across images using min-max scaling. Shaded areas indicate 95% confidence intervals. Images at each rostrocaudal level were warped to a reference section. Abbreviated terms are explained in the list of abbreviations.

### Online repository

Data such as histological images and additional delineations are available upon request and a release is planned in a publicly available data repository.

### Statistical analyses

Custom made scripts in Matlab and Python were used for data analysis and quantification. Fluorescence profiles were analyzed as previously described, using basic programming functions to display mean values with a 95% confidence interval. In the rabies tracing dataset, percentage values were measured per animal; input cells within the CC were not included in these calculations. Mean percentage values were measured across animals, for each individual region, and represented including the standard error of the mean.

## List of anatomical abbreviations

**Prefrontal cortex**

rIns: Insular cortex, rostral part
LO: Lateral orbital cortex
VLO: Ventrolateral orbital cortex
VO: Ventral orbital cortex
MO: Medial orbital cortex
PL: Prelimbic cortex
IL: Infralimbic cortex
DPed: Dorsal peduncular area

**Cingulate cortex**

dACC: Anterior cingulate cortex, dorsal area
vACC: Anterior cingulate cortex, ventral area
RSC: Retrosplenial cortex

**Parahippocampal region**

LEC: Lateral entorhinal cortex
MEC: Medial entorhinal cortex
PER: Perirhinal cortex
POR: Postrhinal cortex
PrS: Presubiculum
PaS: Parasubiculum

**Hippocampal formation**

CA1-3: Cornu ammonis 1-3
DG: Dentate gyrus
Sub: Subiculum

**Sensorimotor cortex**

SSc: Somatosensory cortex
Aud c: Auditory cortex
Vis v: Visual cortex
M1: Primary motor cortex
rM2: Secondary motor cortex, rostral part
cM2: Secondary motor cortex, caudal part

**Other cortices**

cIns: Insular cortex, caudal part
TeA: Temporal association área
Par c: Parietal cortex

**Basal ganglia**

CPu: Caudoputamen
GP: Globus pallidus
VP: Ventral pallidum

**Basal forebrain**

NAc: Nucleus accumbens
NDB: Nucleus of the diagonal band
SIB: Substantia innominata

**Thalamus**

Th MNG: Thalamic midline nuclear group
Th ANG: Thalamic anterior nuclear group
Th LNG: Thalamic lateral nuclear group
Th VNG: Thalamic ventral nuclear group
Th IL: Thalamic intralaminar nuclear group
Th MD: Thalamic mediodorsal nucleus
Meta Th: Metathalamus
Hb: Habenula

**Hypothalamus**

LH: Lateral hypothalamus
MH: Medial hypothalamus
AH: Anterior hypothalamus
PH: Posterior hypothalamus

**Other subcortical areas**

Sept: Septal complex
BNST: Bed nucleus of the stria terminalis
P.Opt: Preoptic area
STh: Subthalamic nucleus
ZI: Zona incerta
Mam.n: Mammillary nucleus

**Piriform area**

rPir: Piriform cortex, rostral area
cPir: Piriform cortex, caudal area
DPir: Deep piriform area
IEn: Intermediate endopiriform nucleus
VEn: Ventral endopiriform nucleus

**Anterior olfactory area**

AOL: Anterior olfactory nucleus, lateral part
AOM: Anterior olfactory nucleus, medial part
AOD: Anterior olfactory nucleus, dorsal part
AOV/P: Anterior olfactory nucleus, ventral/posterior part

**Other olfactory areas**

VTT: Ventral tenia tecta
DTT: Dorsal tenia tecta
Tu: Olfactory tubercle
LOT: Nucleus of the lateral olfactory tract
OB: Olfactory bulb

**Cortical amygdala**

AA: Anterior amygdaloid area
CxA: Cortex amygdala transition zone
ACo: Anterior cortical amygdala
PCo: Posterior cortical amygdala
APir: Amygdala piriform transition zone
IPAC: Interstitial nucleus of the posterior limb of the anterior commissure

**Basolateral complex**

BLA: Basolateral amygdala, anterior
BLP: Basolateral amygdala, posterior
BLV: Basolateral amygdala, ventral
LA: Lateral amygdala
BMA: Basomedial amygdala

**Centromedial amygdala**

CeA: Central amygdala
Med A: Medial amygdala

**Brainstem**

PAG: Periaqueductal grey
SN: Substantia nigra
VTA: Ventral tegmental area
Rn: Raphe nucleus
Rt: Reticular formation
Pn: Pontine nuclei
Other BS: Other brain stem areas

## Results

### Fiber- and cyto-architecture of the claustrum complex

The combined immunohistochemical (IHC) data from PV, MBP, and CB labeling revealed fiber-architectural patterns in the CC suggesting the presence of four distinct domains (Figure 1A-C). Patterns in MBP labeling indicated that the CL can be divided into a dorsal and ventral subregion (dCL and vCL respectively). The PV and CB patterns pointed more to a center-surround organization of the CL, which also helped to define its perimeter. The fiber-architectural patterns for all three markers led us to divide DEn into a medial and lateral subunit (mDEn and lDEn respectively).

In ventral parts of the CL, MBP staining revealed an area with reduced density of myelinated fibers, distinct both from a more dorsal patch of dense labeling in CL and from a dense plexus in layers 5 and 6 of insular cortex (Figure 1A). Dorsal to this ‘MBP-gap’, myelinated fibers extended from the external capsule, moving diagonally towards the insular cortex. The transition from these diagonal fibers to the MBP-gap was used as the main indicator of the dCL-vCL border. Interestingly, the MBP-gap did not fully align with the dense PV-plexus, but only matched with a ventral part of it. However, dorsal parts of the PV-plexus aligned with the patch of MBP-labelled fibers extending diagonally from the external capsule. The dense PV-plexus in the CL contrasted with an absence of labeling in L6 of the insular cortex and in the mDEn (Figure 1B). In general, this plexus provided a good strategy for defining the borders of the CL with the overlying cortex and the DEn. However, in each brain we observed variations in how far the PV-plexus extended dorsoventrally within the CL, and some sections had a PV-plexus that mainly occupied a central part of the CL. The combination of PV and MBP labeling was used to define the border between the CL and insular cortex.

CB-labelled processes formed a ring-like plexus in the CL, with a sparsely labeled central gap (Figure 1C), though a few sections in each brain would not show this in a clear way. Additionally, CB-staining labelled a laminar plexus in layer 6 of insular cortex, which appeared distinct from the CL at central and caudal levels, but less so at rostral levels. The CL-insular border was also visible in CB-labeling, albeit not as clearly as in the other markers. Note that although the PV-plexus, MBP-gap and CB-gap appeared similar in shape, they were spatially misaligned within the CL (Figure 1D-E). The PV plexus extended dorsally past the MBP-gap, occupying both the dCL and the vCL, while the CB-gap aligned with a central region of the PV-plexus and was located dorsally in the vCL (Figure 1E). This is relevant when using only one of these markers to describe experimental data on the CL.

In the DEn, the division into a lateral and medial domain was indicated by all three markers. This was most easily seen in the case of CB staining where the lDEn showed dense immunoreactivity, both resulting from somatic and neurite labeling; CB-labeling in mDEn was considerably sparser. With regards to MBP staining, the mDEn displayed a characteristic striped pattern, separating it from the homogenous labeling seen in the lDEn. The PV labeling was the least discriminative with only the lDEn displaying sparse PV-labeling, which was also not clearly visible in all the material we assessed.

The consistency of our fiber-architectural borders was subsequently assessed along the rostrocaudal axis and across animals by measuring average fluorescence intensity profiles of PV, MBP and CB immunoreactivity (Figure 1F). Brain sections at the rostral, central, and caudal landmarks were selected from each animal, and aligned to a reference section using the image warping pipeline described in the methods section. Profiles from each animal represent average intensity levels of fluorescence as measured in a rectangular selection drawn across the region of interest. The combined profiles for all markers corroborate the differentiation between lDEn and mDEn, mentioned above. They further revealed an overall colocalization of the PV-plexus, MBP gap, and the smaller CB-gap in the CL throughout the rostrocaudal extent of the CL, with the exception of the rostral level, were a slight misalignment of the MBP-gap and PV-plexus was apparent, as described above.

We next aimed to address the unresolved debate on how far CL extends rostrally, by analyzing a series of closely spaced sections. We found that the CB-gap provided the clearest landmark to localize the rostral-most parts of the CL (Figure 2A-B). By following the position of the CB-gap gradually from the rostral landmark (B+0.97) we found that it was clearly visible until about +1.93 mm relative to bregma, but not at B+1.97. The CB-gap aligned consistently with the PV-plexus (Figure 2C-D). However, the PV-plexus in far rostral sections became less distinct due to the prevalence of PV-labeling in surrounding areas. MBP labeling showed little or no indication of the rostral-most parts of the CL (Figure 2E-F). Even though the combination of CB and PV turned out to be good markers for localizing the rostral-most parts of the CL, this combination did not provide a clear border between dorsal and ventral parts of the CL. Therefore, we opted to not subdivide the CL rostral to B+0.97. At levels rostral to B+1.93, the CL could no longer be identified using the current set of criteria, though DEn was still present and could be subdivided into its lateral and medial divisions (Figure 2 all left-hand panels), and this was true even at more rostral levels at B+2.09 (Figure 3B and C – E, left hand panels).

**Figure 2:**
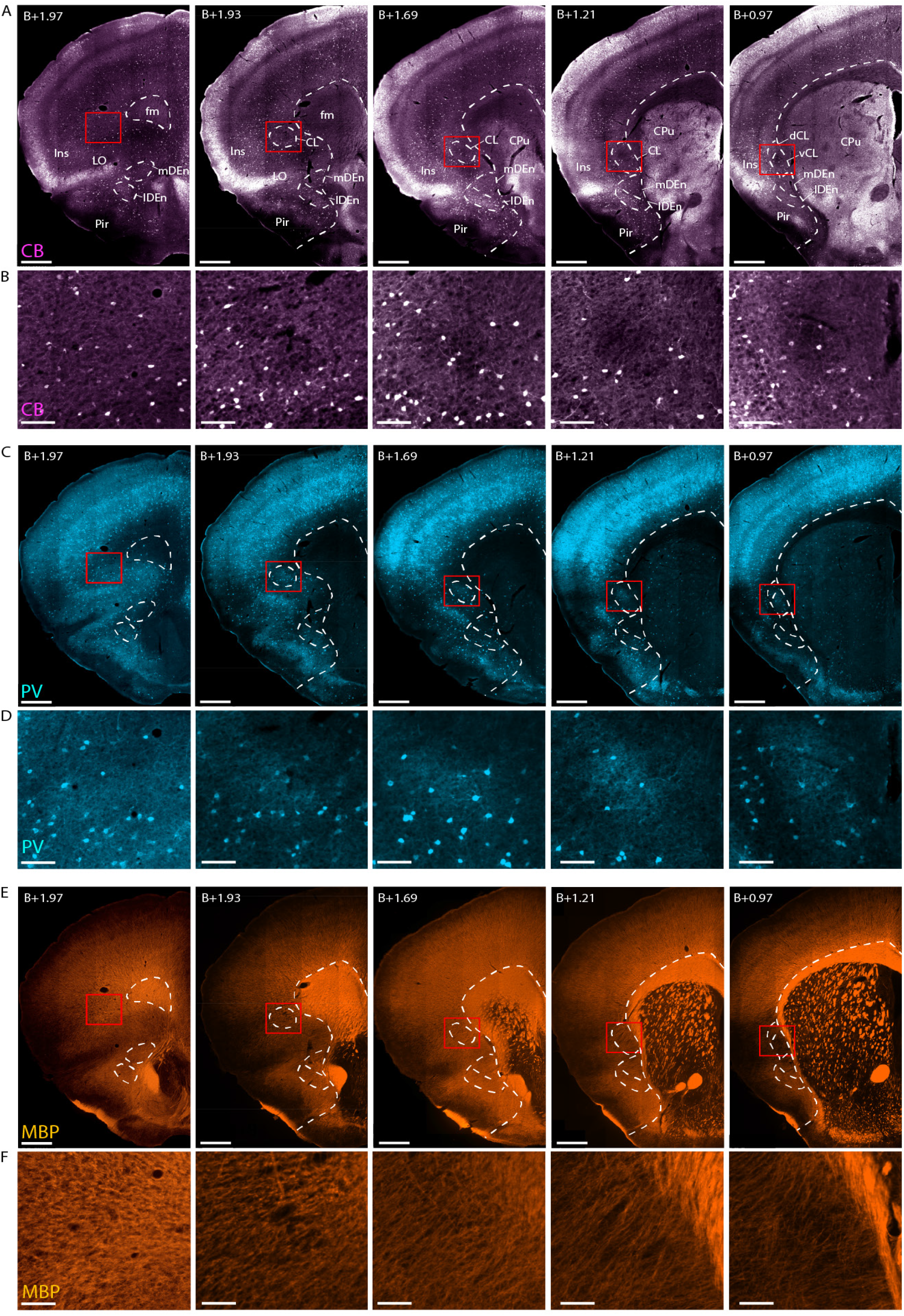
Delineating the rostral-most parts of the claustrum. Immunohistochemical labeling of CB, PV and MBP in five sections, taken at gradually more rostral levels (left to right, indicated by level rostral to Bregma (B)). Red squares indicate the location of each inset. **A-B)** Location of the CB-gap, revealing the CL until B+1.93. **C-D)** Images of the same sections (aligned to the CB-image in A-B) stained for PV. Note the presence of dense PV neuropil aligned with the CB-gap. **E-F)** MBP labeling indicates the position of vCl only in the most posterior of the five sections. Scale bars measure 500μm in A, C, E and 100μm in B, D, F.

**Figure 3:**
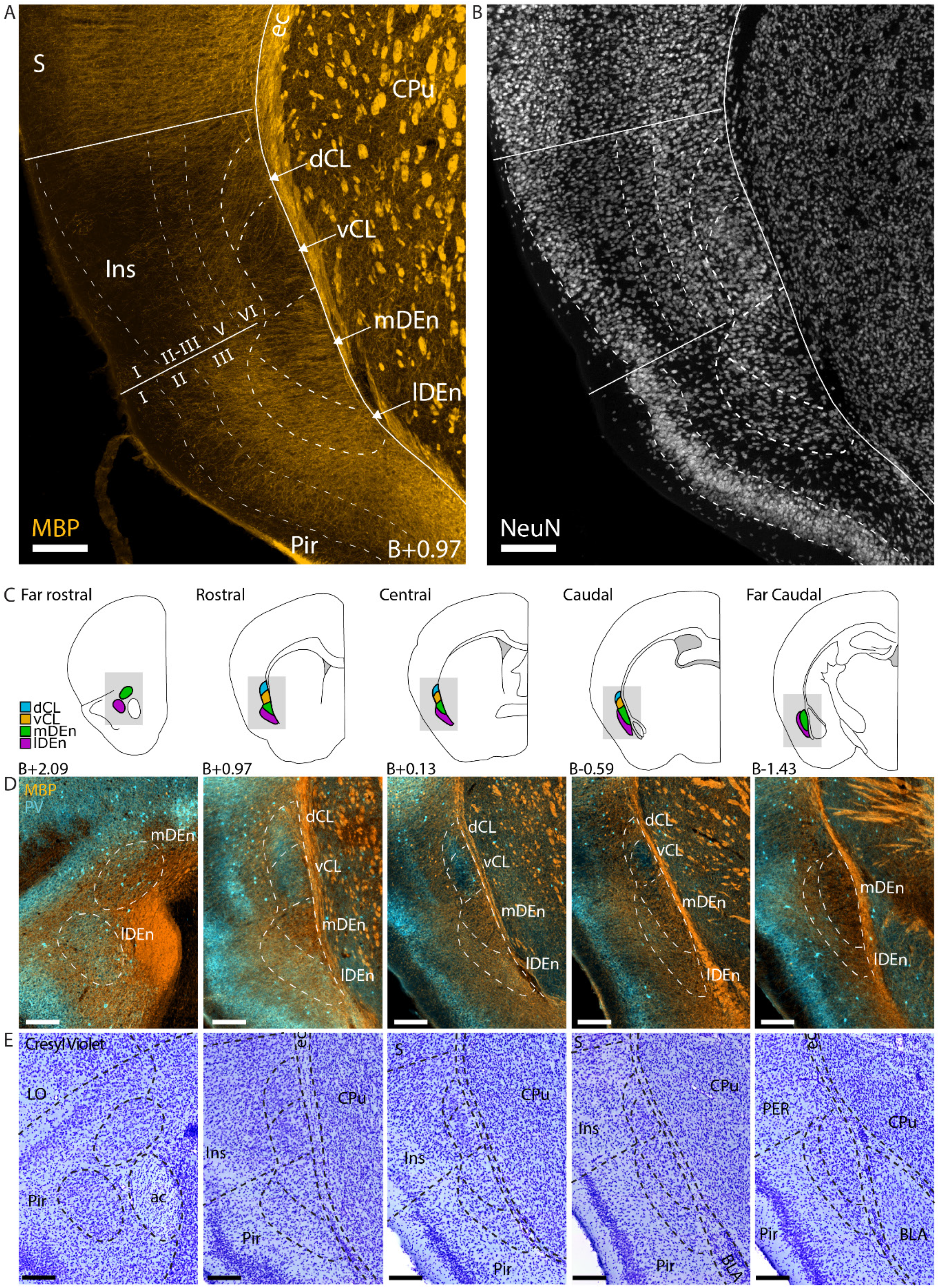
Comparison of fiber- and cyto-architecture of the CC. **A-B)** Immunohistochemical staining against MBP and NeuN. Delineations based on fiber-architectural patterns overlap with those seen in cytoarchitecture. **C)** Schematic representation of the CC at 5 rostrocaudal landmarks. Shaded areas indicate inset location in D-E. Approximate location to bregma is shown at each landmark. **D)** Co-expression of MBP and PV at each landmark. **E)** Cresyl Violet staining of the same sections shown in D. Scale bars measure 200μm. Abbreviated terms are explained in the list of abbreviations.

To compare fiber- and cyto-architecture in the CC, we did IHC staining against PV and MBP in addition to either NeuN or Cresyl Violet (Figure 3). Together, these experiments revealed distinct cyto-architectural features within the fiber-based subregions of the CC (Figure 3A-B). We observed that the cell-arrangement in the CC was often aligned with the direction of MBP-labelled fibers. Diagonal columns of cells could be seen in dCL, following the fibers that extended from the external capsule towards insular cortex, whereas cells in vCL were arranged in a circular structure, matching the shape of the MBP gap. The mDEn showed a columnar arrangement of cells that were aligned to the stripes in MBP labeling. In comparison, the lDEn had a more laminar arrangement of cells, following the piriform cortex. These features were generally clearer in Cresyl Violet staining than NeuN labeling.

In a final series of experiments, we did immunostaining against PV and MBP followed by Nissl staining in the same tissue, to corroborate our definitions of the most rostral areas of CC and also to precisely define its caudal-most parts. We also aimed to increase our resolution along the rostrocaudal axis, so we stained and mounted every second coronal section, allowing us to study the gradual change of claustral borders with close rostrocaudal increments (80μm between sections; Figure 3C-E). With this approach we delineated the CC at far rostral and far caudal levels. The set of images used to delineate the entire rostrocaudal length of the CC will be uploaded to the online repository.

At far rostral levels (B+2.09), the mDEn appeared in the medial most parts of the piriform cortex, dorsal to the anterior commissure; the lDEn was distinct from the mDEn at this level and was positioned deep to piriform cortex on the lateral side of the anterior commissure (Figure 3D-E, left-most panels). MBP labelled fibers were sparser in far rostral lDEn than in surrounding areas. In particular, the area between lDEn and mDEn was densely stained with MBP. Far rostral mDEn was also indicated by MBP labeling but this was not always easy to see. In some brains the far rostral lDEn had a clear PV plexus. Cyto-architecturally, both areas were more densely populated than the surrounding parts of piriform cortex. The CL was not visible at far rostral levels, but as shown in figure 2, it followed the dorsoventral position of the insular cortex and was distinct from DEn in sections at levels where it was possible to differentiate the lateral orbital cortex.

At far caudal levels (B-1.43), the lDEn covered the entire lateral border of the mDEn (Figure 3D-E, rightmost panel). The two regions were distinguished by sparse labeling of MBP in the mDEn compared to lDEn. The mDEn was absent of PV labeling, while the lDEn showed some labeling. Cyto-architecturally, the far caudal lDEn had a laminar appearance, while the mDEn appeared more irregular. Both regions extended as far caudal as the piriform cortex, which was gradually replaced by the lateral entorhinal cortex (LEC). The transition from lDEn and mDEn to L6 of LEC was easier to identify in MBP and PV staining than with cyto-architecture.

### Gene expression patterns corroborate architectonic delineations

To expand upon the current toolbox of genetic markers for the CC, we used a list of genes acquired from a chromatin immunoprecipitation sequencing (ChIP-Seq) analysis of micro dissected claustrum tissue (unpublished own material). We then screened the Allen *in situ* hybridization (ISH) database for these genes, in addition to using the in-built differential gene search in the Allen ISH database for candidate markers in the claustrum, endopiriform nucleus, insular and piriform cortices (© 2006 Allen Institute for Brain Science, ISH Data, Available from: https://mouse.brain-map.org/). Finally, we searched the literature for genetic markers used to identify deep cortical layers. From more than one thousand ISH experiments we screened a short list of eligible candidates as presented in Table 3.

As several genes showed similar distribution patterns, we selected a few to analyze in more detail (Figure 4A). For these genes, we downloaded images of coronal slices at the five rostrocaudal landmarks and aligned them to a reference brain section using the image warping pipeline described in the methods (Figure 4B-E). We did not delineate the CC during the image warping procedure but used well-defined borders of surrounding areas like the external capsule and borders between insular, piriform and somatosensory cortex. Delineations were drawn on the reference section, which was labelled with MBP, PV and Cresyl Violet, without viewing the gene expression. PV-labeling was selected for the background as this provided the clearest visual distinction to the superimposed data.

**Figure 4.**
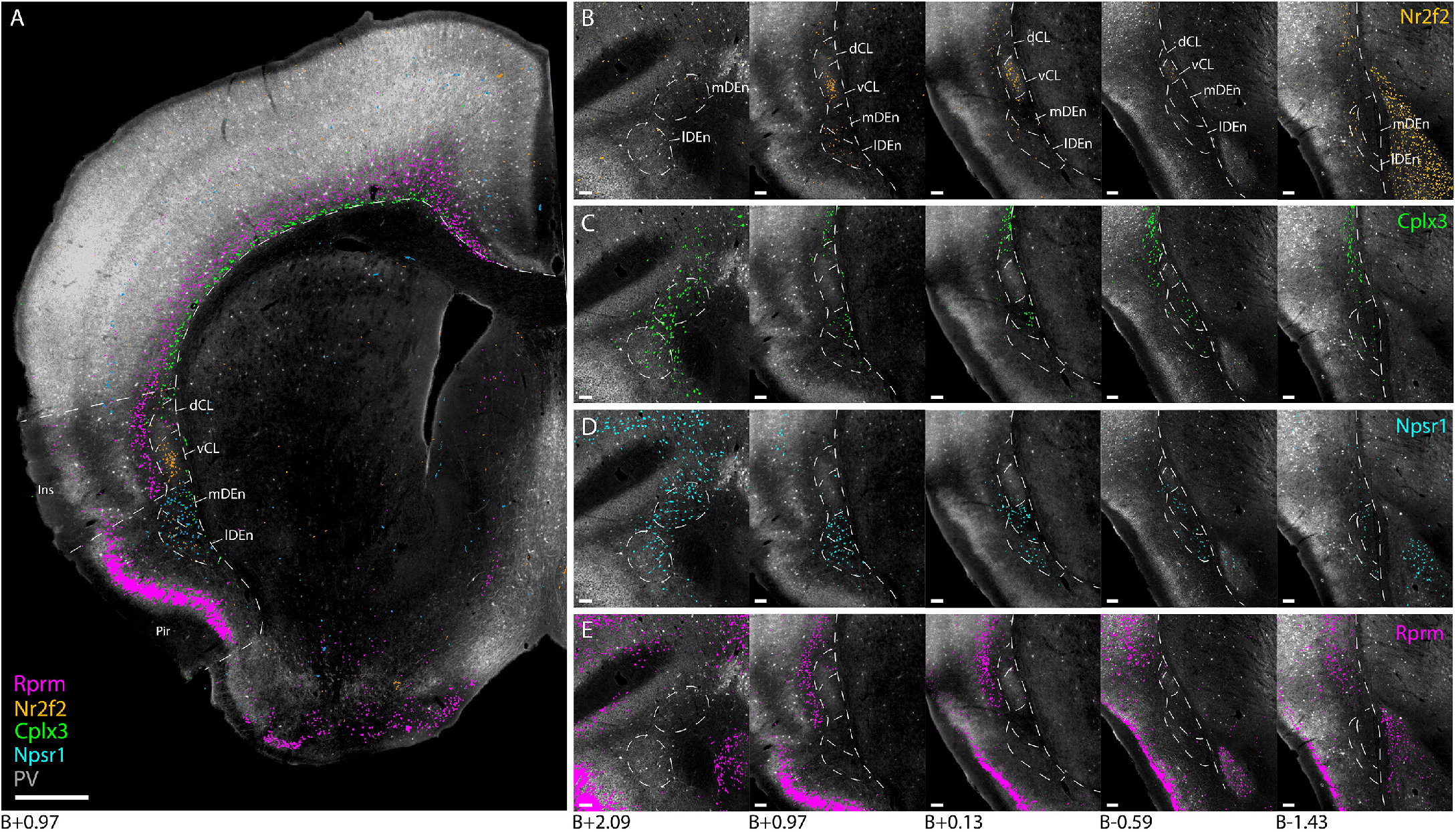
Genetic marker expression in the claustrum complex. Gene expression images were collected from the Allen ISH database and aligned to a reference section (see method section for details). **A)** Combined expression of *Nr2f2, Cplx3, Npsr1* and *Rprm* genes superimposed onto a reference section located at the rostral landmark. **B-E)** Individual gene expression in the claustrum region from the five rostrocaudal levels taken at the previously defined landmarks (see methods). Scale bars measure 500μm in A, and 100μm in B-E.

We found similar expression patterns among some of the genetic markers. Genes such as *Nr2f2* and *Rxfp1*, displayed a dense, centered expression within vCL, in addition to sparser labeling in lDEn (Figure 4B), and genes like *Cplx3, Ctgf* and *Galnt10* showed confined expression along the cortical subplate that stopped upon reaching the CL but reappeared in mDEn (Figure 4C). The *Npsr1* gene had dense and highly specific expression in both mDEn and lDEn (Figure 4D). Among layer 6 markers, the *Rprm* gene stood out with a dense laminar expression throughout L6 of the entire neocortex, and laterally along the outside of the CC (Figure 4E). The expression of the *Rprm* gene was clearly aligned with the area lacking PV labeling that was used to delineate layer 6 of insular cortex. Note that some of these markers, (*Nr2f2, Cplx3* and *Ctgf*) have been characterized in previous studies on gene expression in the CL (Bruguier et al., 2020; Erwin et al., 2021; Wang et al., 2017), but not in comparison to a fiber-architectural reference. To our knowledge, the *Npsr1* and *Rprm* genes have not been characterized before for the CC.

### Fiber-architectural location of the retrosplenial-projecting claustrum pocket

A dense pocket of cells can be labelled in the CL by injecting a retrograde tracer into the retrosplenial cortex (RSC; Zingg et al., 2018). To assess the location of this RSC-projecting pocket of CL cells (CL_RSC_ -pocket), relative to our fiber-architectural markers, we injected cholera toxin subunit B (CTB) into the RSC of C57BL6J mice (n=6). The tissue was then stained for the expression of MBP and PV (Figure 5A-B). The dense CL_RSC_ -pocket of labelled neurons shows a strong overlap with the MBP-gap, though a more dispersed population of cells was seen in surrounding parts of the CL. Fluorescence profiles, measured along the mediolateral and dorsoventral axes through the center of the CL_RSC_ -pocket, showed a good alignment of the labelled neurons along both axes with the peak of PV fluorescence and trough of MBP fluorescence (Figure 5C). Note that at rostral levels the MBP trough was shifted slightly ventral to the CLRSC-pocket.

**Figure 5:**
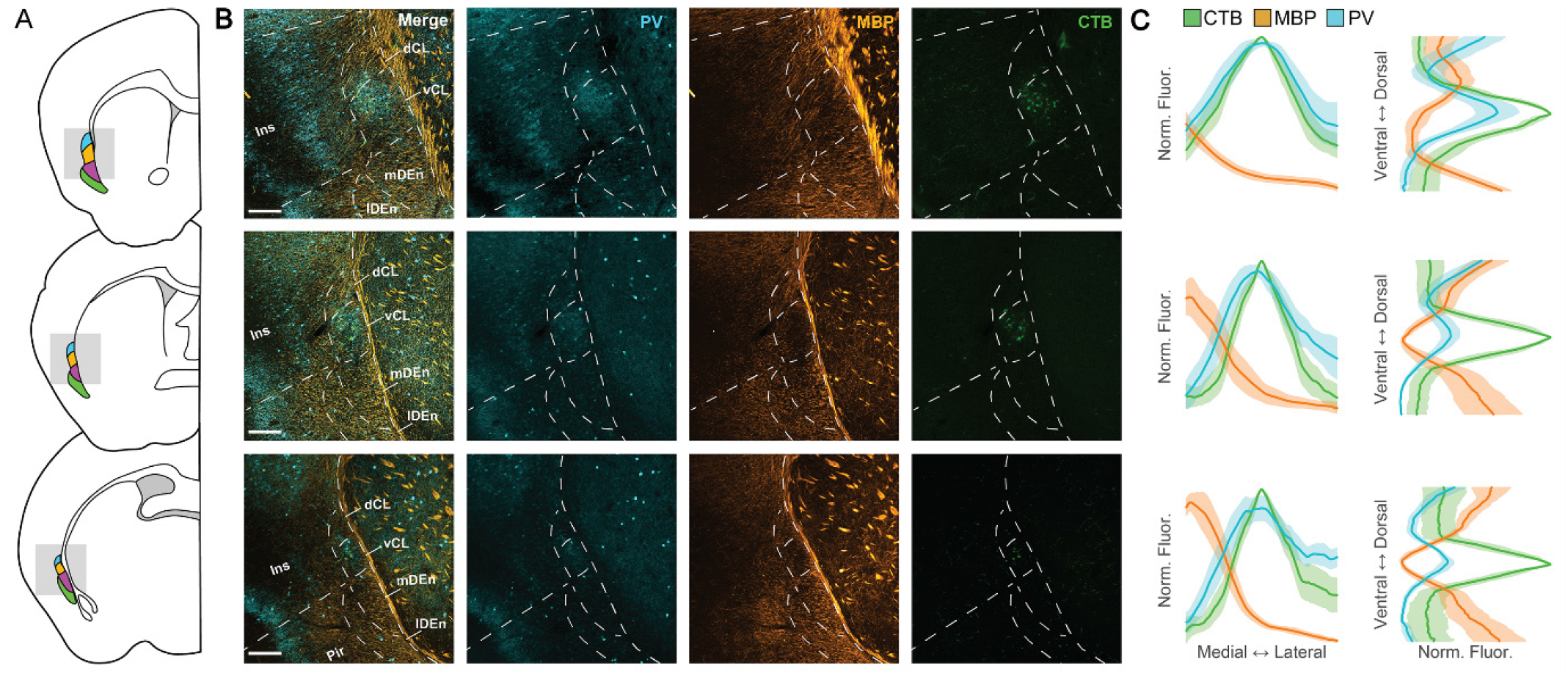
RSC-projecting claustrum neurons overlap with the PV peak and the MBP trough in the CL. **A)** Schematic representation of tissue sections used for IHC and further analysis. **B)** Representative confocal images of claustra from the sections in A with retrogradely labeled claustrum neurons (green) and immuno-staining against MBP (orange) and PV (cyan). Scale bars represent 200 μm. **C)** Normalized fluorescence traces for each of the sections in A and B along the mediolateral (left) and dorsoventral (right) axes. Shaded areas represent 95% confidence intervals.

### Brain-wide monosynaptic inputs to the mouse CC

Using the *CC-EDGE::Tre-Tight-THAG* transgenic mouse line, we conducted monosynaptic rabies-tracing of brain-wide inputs to all subregions of the CC. The location of each input and starter cell was determined based on Cresyl Violet staining. As such, the precise location of every cell could be determined within each animal. Input cells were found in a myriad of cortical and subcortical areas, expanding the input connectivity known from the literature (Figure 6). Substantial inputs were identified coming from known input areas like the anterior cingulate cortex (Figure 6A-B), the amygdala (Figure 6C-D) and anterior olfactory areas (Figure 6E-F), and from less documented input areas like the hippocampal formation (Figure 6G-H).

**Figure 6:**
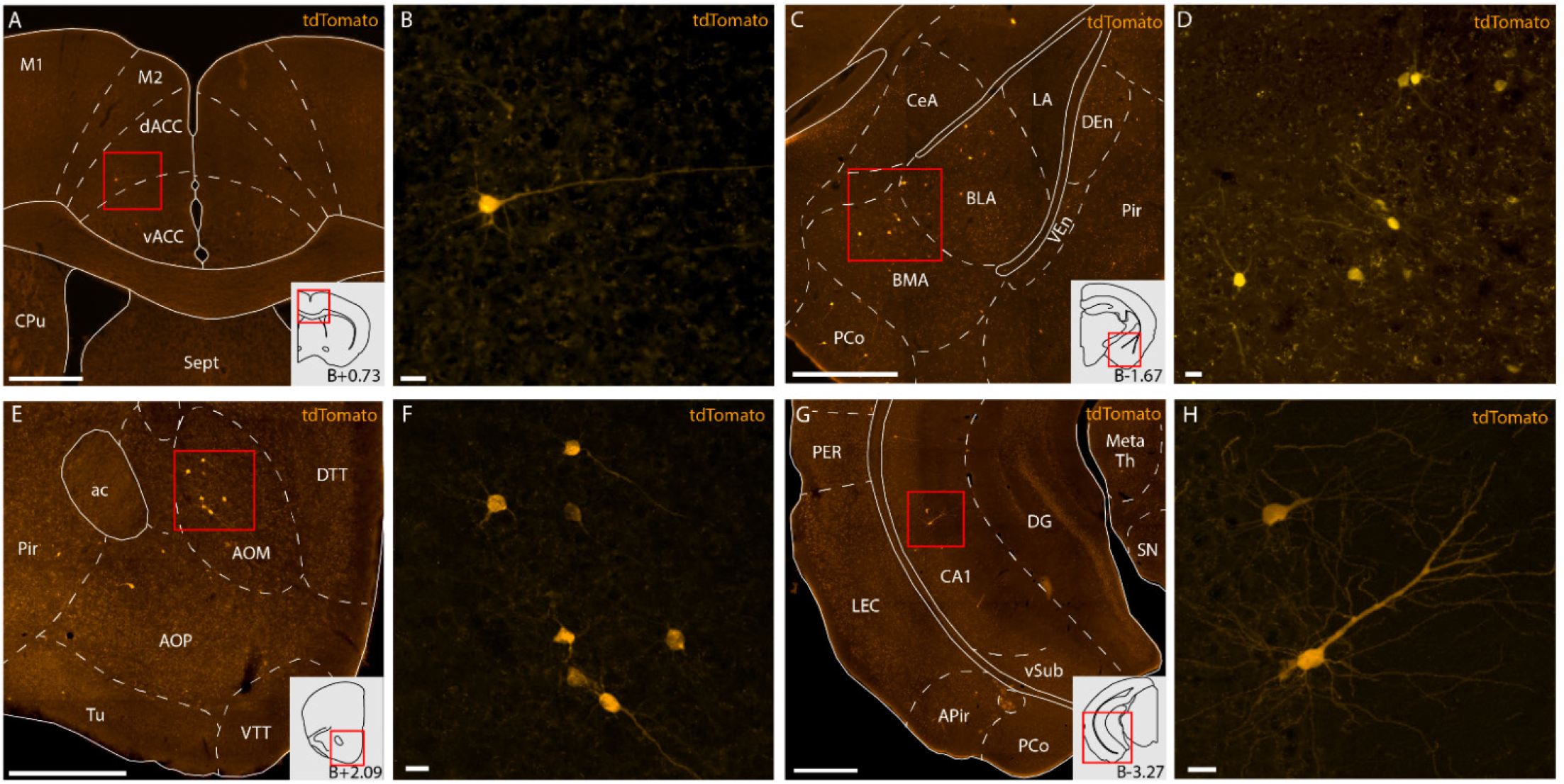
Example sections from rabies tracing dataset showcasing monosynaptic input cells projecting to the claustrum complex (CC) from anterior cingulate cortex (ACC; A-B), basolateral amygdala (BLA; C-D), medial anterior olfactory area (AOM; E-F) and cornu ammonis 1 (CA1; G-H). All delineations are based on Cresyl Violet staining of the same tissue. Input cells are labelled by a tdTomato-tag expressed by the rabies virus. Schematics in the bottom right corner of A, D, E and G show outlines of each representative coronal section, and the approximate distance relative to bregma. Scale bars measure 500μm in A, C, E, G and 20μm in B,D,F,H. Abbreviated terms are explained in the list of abbreviations.

Starter cells were immunohistochemically identified by the co-expression of the 2A linker protein and a tdTomato tag (Figure 7A); input cells were identified by the expression of tdTomato, but not the 2A linker protein. We observed no co-expression in double *in situ* staining against the THAG sequence, found in transgene expressing cells of the *CC-EDGE::Tre-Tight-THAG* line, and GAD67, a general inhibitory marker, indicating that the starter cells were excitatory neurons (Figure 7B). Across 13 animals, 95.8% (±1.5% SE) of all starter cells were located within the CC; a few were found in nearby cortices (Table 4). Within the CC, starter cells were found in each dorsoventral subregion, although the dCL only contained a minority (Figure 7C). The starter cell population covered most of the rostrocaudal extent of the CC, the exception being far caudal parts of the mDEn and lDEn.

**Table 4:**
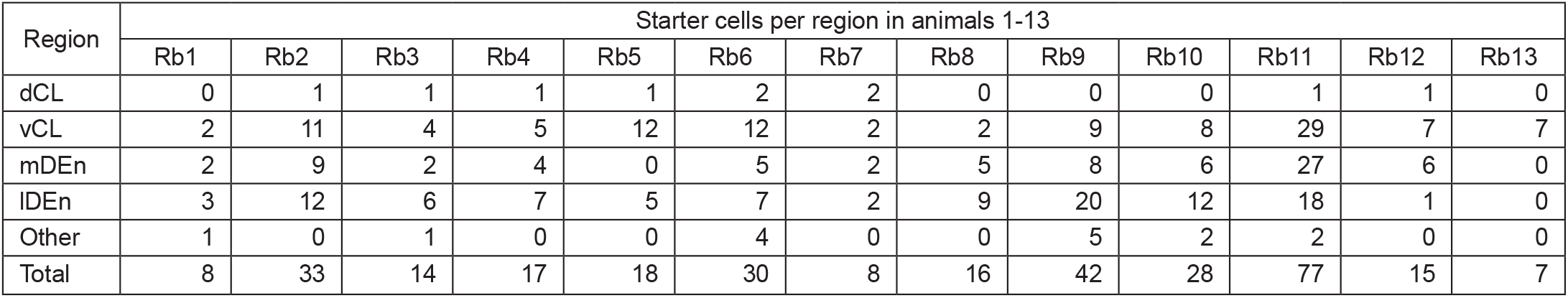
Starter cell distributions

| Region | Starter cells per region in animals 1-13 |  |  |  |  |  |  |  |  |  |  |  |  |
| --- | --- | --- | --- | --- | --- | --- | --- | --- | --- | --- | --- | --- | --- |
|  | Rb1 | Rb2 | Rb3 | Rb4 | Rb5 | Rb6 | Rb7 | Rb8 | Rb9 | Rb10 | Rb11 | Rb12 | Rb13 |
| dCL | 0 | 1 | 1 | 1 | 1 | 2 | 2 | 0 | 0 | 0 | 1 | 1 | 0 |
| vCL | 2 | 11 | 4 | 5 | 12 | 12 | 2 | 2 | 9 | 8 | 29 | 7 | 7 |
| mDEn | 2 | 9 | 2 | 4 | 0 | 5 | 2 | 5 | 8 | 6 | 27 | 6 | 0 |
| IDEn | 3 | 12 | 6 | 7 | 5 | 7 | 2 | 9 | 20 | 12 | 18 | 1 | 0 |
| Other | 1 | 0 | 1 | 0 | 0 | 4 | 0 | 0 | 5 | 2 | 2 | 0 | 0 |
| Total | 8 | 33 | 14 | 17 | 18 | 30 | 8 | 16 | 42 | 28 | 77 | 15 | 7 |

**Table 7:**
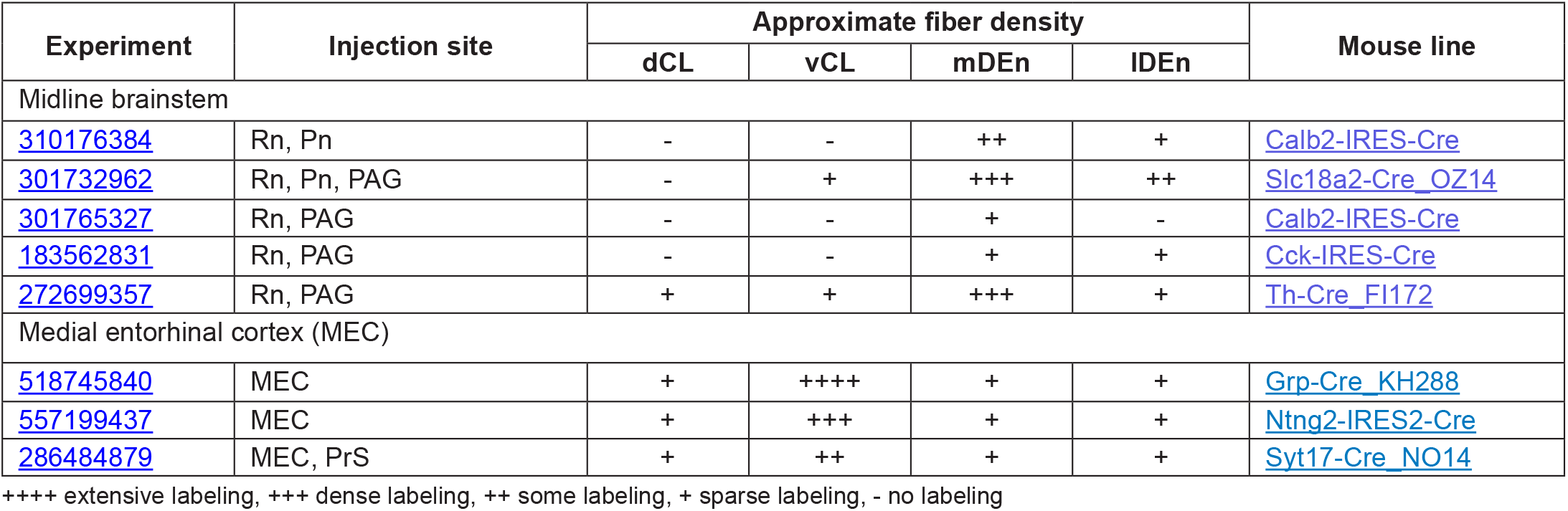
List of experiments collected from the Allen Projection database

**Figure 7:**
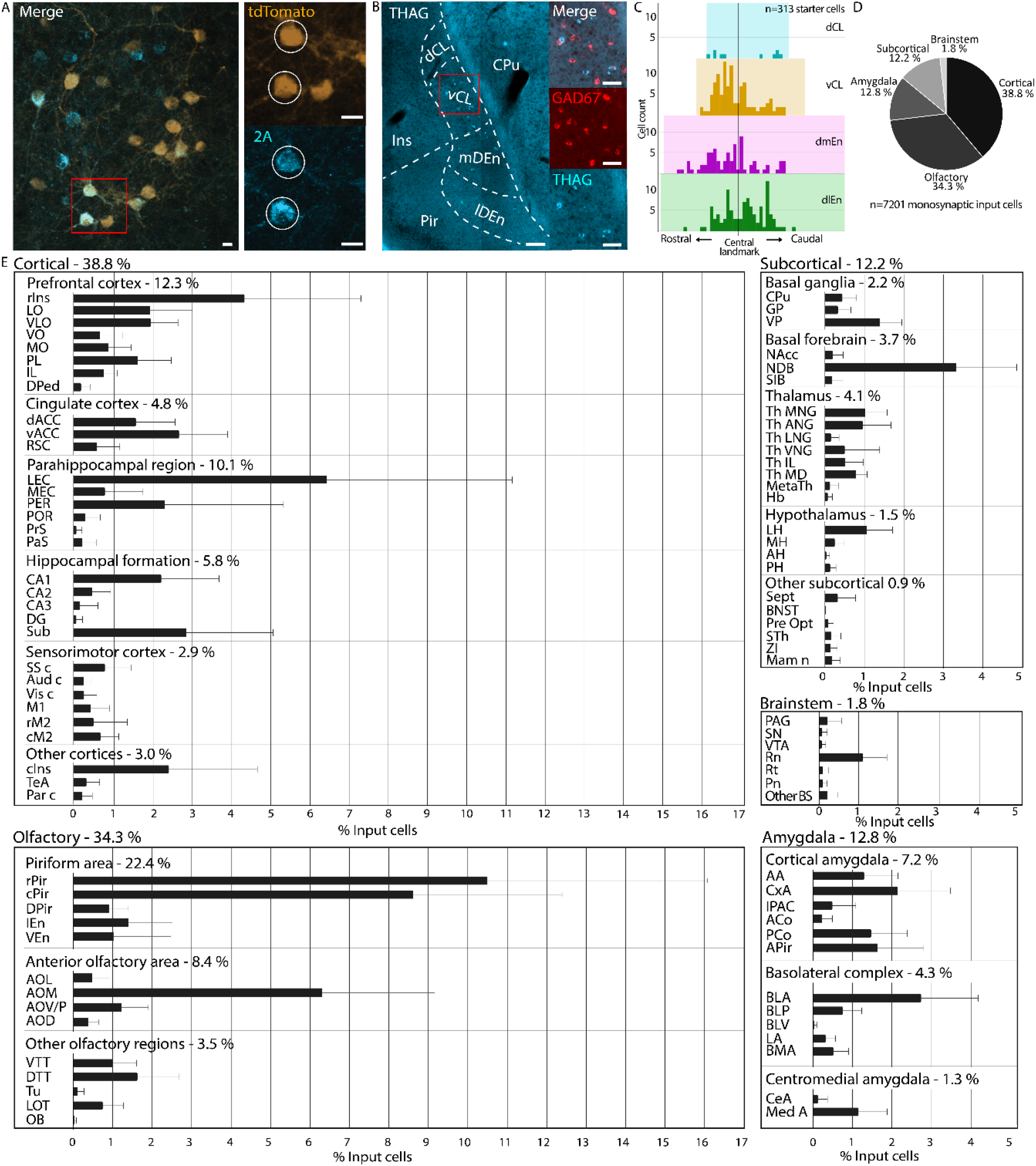
Brain-wide monosynaptic input tracing to the claustrum complex. **A)** Confocal image of starter cells in the CL, identified by the co-expression of tdTomato, expressed by the rabies virus, and the 2A linker protein, expressed in transgene expressing cells. Scale bars measure 20 μm. **B)** Double *in situ* hybridization against THAG sequence, found in transgene expressing cells, and the general inhibitory marker GAD67. **C)** Histograms showing rostrocaudal position of pooled starter cells from all animals within claustral subregions. **D)** Coarse overview of monosynaptic inputs to the CC. **E)** Detailed overview of monosynaptic inputs to the CC (n=7201 cells, error bars show the standard error of the mean). Abbreviated terms are explained in the list of abbreviations.

We divided the input connectivity into 5 major categories (Figure 7D): cortical (38.8±3.6% SE), subcortical (12.2±0.7% SE), olfactory (34.3±3.4% SE), amygdala (12.8±1.3% SE) and brainstem (1.8±0.3% SE). In total, we found inputs in 89 cyto-architecturally defined input regions (Figure 7E). This included known input areas to the CL like the prefrontal cortex, thalamic nuclei, and the basolateral amygdala. Known inputs to the DEn like anterior olfactory areas, cortical amygdala and lateral entorhinal cortex were also prevalent in the dataset. Additionally, we found considerable inputs from hippocampal regions, mainly in cornu ammonis 1 (CA1). There were also cells in CA2, CA3 and the dentate gyrus (DG), which have not been described before. Other yet undescribed input regions include the habenula (Hb), bed nucleus of the stria terminalis (BNST) and zona incerta (ZI) (Table 7-1).

A small fraction of input cells was found in the contralateral hemisphere, and these cells were largely present in prefrontal and cingulate areas (Table 7-2). Some regions like the anterior cingulate (ACC) and prelimbic cortex (PL), showed a balanced distribution of ipsi- and contralateral inputs, whereas areas like the basolateral amygdala (BLA) and lateral entorhinal cortex (LEC) showed a skewed distribution with only a small fraction of contralateral inputs. Notably, we observed contralateral inputs from the nucleus of the diagonal band (NDB), lateral hypothalamus (LH) and the nucleus of the lateral olfactory tract (LOT), which have not been previously shown to project bilaterally to the CC. Areas along the midline of the brain were not categorized as ipsi- or contralateral.

We selected a few representative brains with decreasing proportions of starter cells within the CL (Figure 8A). Among these, the ones with more CL-starter cells had a higher representation of cortical inputs, and less olfactory and brainstem inputs (Figure 8B). Inputs from medial entorhinal cortex (MEC) and CA1 were more prevalent in brains with a high percentage of CL starter cells, while raphe nuclei (Rn) and lateral anterior olfactory area (AOL) inputs showed an opposite trend (Figure 8C). Using the Allen Projection database, we found a corresponding pattern in afferent projections from MEC and midline regions of the brainstem (Table 5). Images collected from these databases were fitted to our reference brain using image warping, allowing the signal to be superimposed onto sections delineated based on fiber- and cyto-architecture. Corroborating the input patterns seen in our dataset, the MEC injected brains showed dense labeling in the vCL (Figure 8D), while injections in the midline of the brainstem showed labeling surrounding the vCL, predominantly located in the mDEn (Figure 8E).

**Figure 8:**
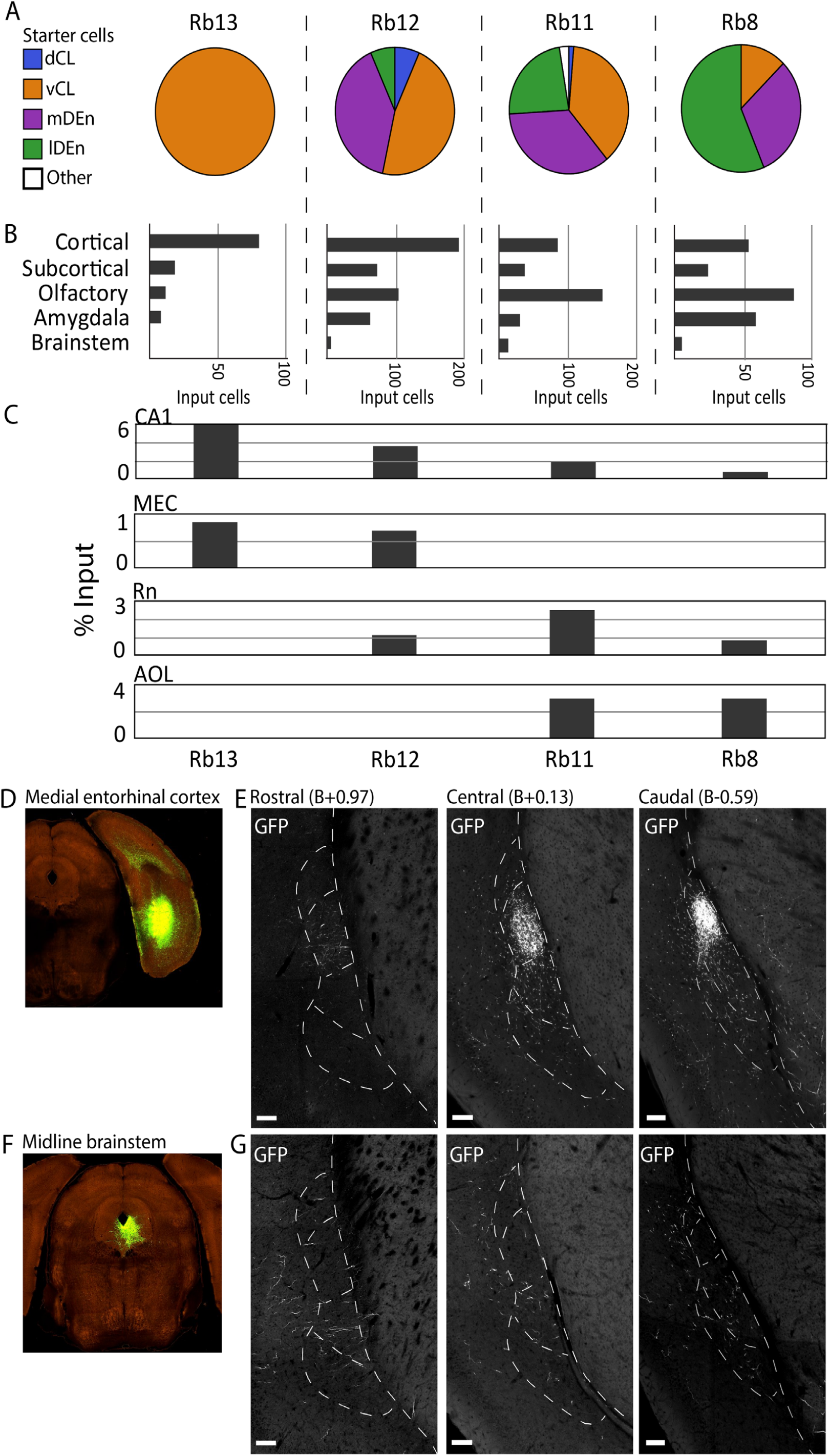
Comparison of input and starter cell populations. **A)** Starter cell distributions in four representative brains displaying various proportions of starter cells in the CC. **B)** Coarse input distributions in the same four brains. **C)** Percentage of inputs from CA1, medial entorhinal cortex (MEC), raphe nuclei (Rn) and lateral anterior olfactory nucleus (AOL), in representative brains. **D)** Injection site for experiment 518745840 with an injection of rAAV-EGFP anterograde tracer in the MEC (© 2017 Allen Institute for Brain Science, Projection Dataset, Available from: https://connectivity.brain-map. org/). **E)** Axonal projections from MEC innervating rostral, central and caudal parts of the claustrum complex. Images were warped onto matching reference-sections to align the signal with fiber- and cyto-architectural delineations. **F)** Same as D, but from experiment 272699357 with the same anterograde tracer deposited in midline brainstem areas Rn and periaqueductal grey. **G)** Same as E, but from experiment 272699357. Scale bars measure 100 μm. Image credit (D-G): Allen Institute.

## Discussion

Borders for the rodent CC are difficult to define, leading to substantial variation in how this brain region is delineated (Bruguier et al., 2020; Dillingham et al., 2019; Fang et al., 2021; Smith et al., 2019; Wang et al., 2022). We combined different strategies in search of overlapping patterns to delineate the CC as well as to define its constituting subdivisions. As a result, we present a multifaceted definition for the borders of the CC, based on the expression patterns of multiple, methodologically different markers, and aided by established anatomical features of adjacent cortices like the PV-labelled neuropil in L5 of insular cortex and the distribution of CB positive cells in L5-6 of neocortex (Alcantara et al., 1993; Hof et al., 1999; Tremblay et al., 2016).

We present a highly detailed characterization of myelinated fiber-patterns in the mouse CC. In general, MBP staining in the CC displayed intricate patterns that were highly useful for delineation. We observed clear differences in the amount of myelination within subregions of the CC, which is also the case in marmosets, where the CL is more myelinated than the DEn (Pham et al., 2019). Our motivation for studying MBP-labeling was also due to the evolutionarily preserved fiber-tracts encapsulating the CC in mammals (Bruguier et al., 2020; Buchanan & Johnson, 2011; Kowianski et al., 1999). As indicated by MBP staining lateral to CL, an extreme-capsule-equivalent could be present in mice, albeit merged into L5-6 of insular cortex. It would be interesting to see if similar myelin patterns exist in other animals that lack a clear extreme capsule, such as rats or fruit bats.

Our research also led to novel findings in the expression of PV and CB in the CC. We discovered a PV-plexus in the lDEn which has not been described in prior studies (Druga et al., 1993; Real et al., 2003; Suzuki & Bekkers, 2010). Additionally, we characterized a ring-like CB-plexus in the CL, which has not been identified before, although sparsity of CB-labeling has been described (Celio, 1990; Davila et al., 2005; Druga et al., 1993). The CB-plexus was also visible in the CL anterior to the striatum, corroborating borders indicated by *Crym* ISH-labeling (Dillingham et al., 2017). The unique appearance of the CB-plexus, which reliably follows the PV-plexus in the CL also at the rostral extremes, makes a solid argument for the existence of a CL-domain anterior to the striatum. Earlier work have placed the rostral-most borders for the CL on the ventromedial end of the forceps minor (Grasby & Talk, 2013; Jankowski & O’Mara, 2015), but this area likely belongs to the cortex as it lacks a clear PV-plexus and genetic markers for the CL (Mathur et al., 2009).

Since CB is expressed both in excitatory and inhibitory cell-types (DeFelipe, 1997; Gonchar & Burkhalter, 1997), it is difficult to make functional assumptions about the CB-plexus in the CL. However, there is a similarity between the CB-plexus and somatostatin (SST) labeling in the CL, which labels a major sub-type of interneurons (Graf et al., 2020; Tremblay et al., 2016). Considering that some CB-labelled cells co-express PV, it could be that the CB-plexus represents an inhibitory network in the CL. Fibers expressing calretinin (CR), a marker that is mainly present in inhibitory neurons and rarely co-expresses with CB, occupy the perimeter of the CL in a similar way to the CB-plexus (Barinka & Druga, 2010; Davila et al., 2005; Real et al., 2003). Together, these networks of CB-, CR- and SST-positive neurites could represent an “inhibitory surround” that regulates the central pocket of excitatory projection neurons in the CL.

Anatomical descriptions of the rodent CL generally fall into two categories: dorsoventral partitioning and center-surround organization. Since we observed indications of both categories in our dataset, these two organizational principles might co-exist (Figure 9). There was a clear dorsoventral variation in the extent of myelination and cellular density in the CL. Conversely, a center-surround organization was seen in the ring-like CB-plexus, and by the CLRSC-pocket, which occupied a central compartment of the PV-plexus. The co-existence of dorsoventral gradients with a center-surround structure, is similar to what is proposed by Marriott et al. (2021). The transition from insular to piriform cortex has traditionally been considered to indicate the border between the CL and DEn, which our results show as well and further provides better foundation for this border. Note that we did not consider the intermediate and ventral endopiriform nuclei to be part of the CC, following ontogenetic differences (Watson & Puelles, 2017). These regions were therefore not characterized in our study.

**Figure 9:**
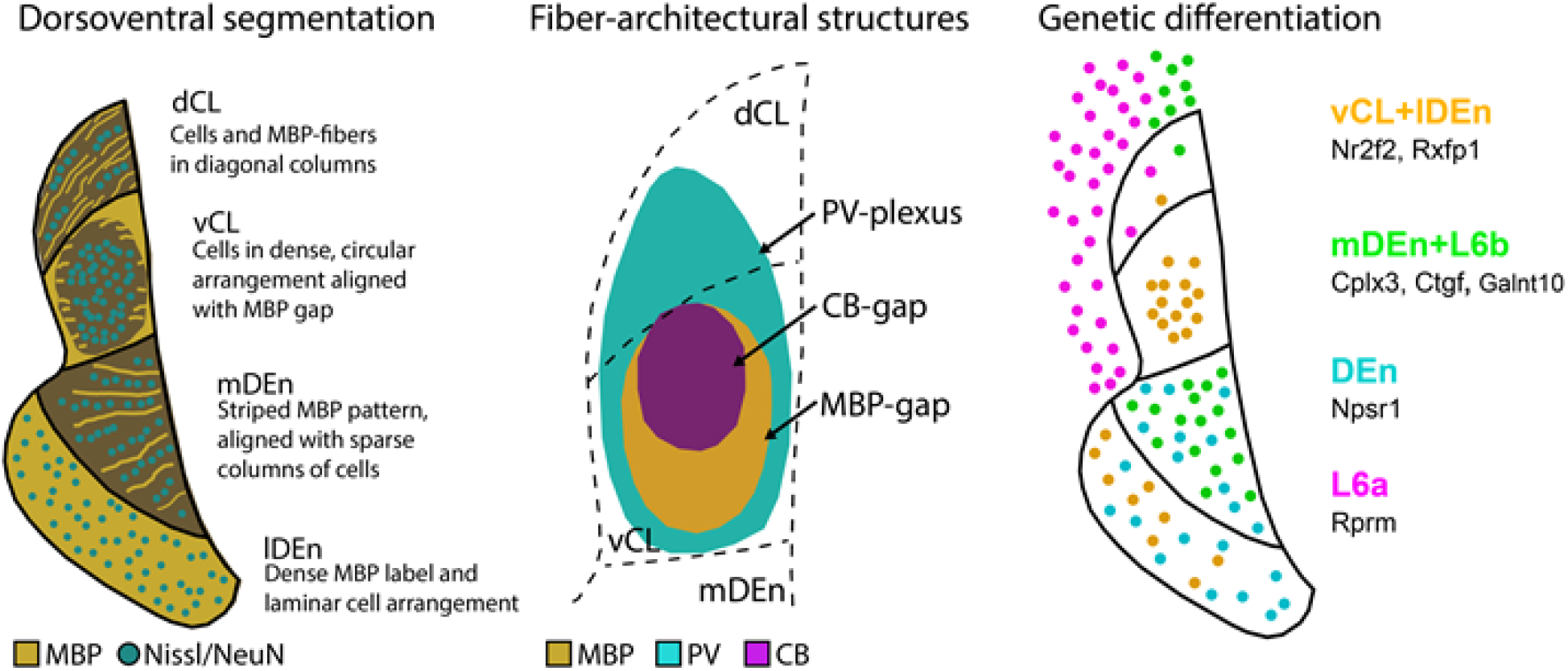
Schematic representation of our characterization of the CC, describing fiber- and cyto-architectural markers and genetic diversity.

Gene expression gives insight into the genetic diversity of brain regions, but only rarely do they express selectively in one area (Lein et al., 2007). For the CC, known markers also express in the adjacent insular cortex (Wang et al, 2017), which is what we observed as well. This is likely associated with the shared developmental origin of these structures from the lateral pallium. During development, cells destined for the CL and DEn migrate alongside those going to the insular cortex, and they are genetically distinguishable by the marker *Nurr1* (Watson & Puelles, 2017). Still, although *Nurr1* is clearly enriched in the CC it also expresses in the insular cortex (Fang et al., 2021). Therefore, the use of genetic markers to differentiate between cortex and CL warrants some caution. Among the myriad of markers screened in this study, we observed two prevalent expression patterns in the CC, where genes like *Nr2f2* labeled the vCL and lDEn, while genes like *Cplx3* labelled the mDEn. This corroborates a genetic diversity previously described in the literature (Erwin et al., 2021; Watson & Puelles, 2017), and together with the unique features seen in our own data, distinguishes the mDEn area from other parts of the CC. Interestingly, projections from medial brain-stem areas like the dorsal raphe nuclei and the periaqueductal grey selectively innervate the mDEn. These regions comprise a substantial fraction of subcortical inputs in our dataset and could be pertinent to the functional properties of the mDEn.

With our multifaceted delineation strategy, we could precisely compare ISH-data to a fiber-architectural reference frame, allowing us to disentangle labelled cells in the CC from the surrounding cortex. Our claustrocortical border, primarily defined by the PV-plexus, was also indicated by the L6 marker *Rprm*. While markers like *Synpr1* and *Nurr1* are valuable tools for locating the CC, they likely exhibit some co-expression in adjacent cortex (Arimatsu et al., 2003), warranting caution for how they are used to define the dorsal border between the CL and insular cortex (Binks et al., 2019; Fang et al., 2021). Furthermore, subdivisions of the DEn based on *Nurr1*-expression (Fang et al., 2021) do not capture the clear mediolateral border we observed in multiple fiber-architectural markers, which highlights the importance of our multifaceted delineation strategy. A crucial element in our approach was the co-expression of multiple markers in the same tissue, which led us to discover that the MBP-gap only occupied a ventral part of the claustral PV-plexus. This forms a solid argument for subdividing the CL into a dorsal and ventral domain, thus opposing a recent delineation where theCL was confined to the MBP-gap and not subdivided into a dorsal and ventral component (Wang et al., 2022).

As part of our characterization, we also conducted a comprehensive tracing study of brain-wide inputs to the CC, using the CC-EDGE transgenic mouse-line (Blankvoort et al., 2018). A unique aspect of our dataset is that the locations of all cells were anchored to cyto-architectonically defined brain areas, instead of mapping them loosely to atlas delineations. In total we categorized inputs from 89 different brain regions, which considerably expands previous tracing studies targeting the CC. Further, we report brain-wide inputs to the DEn, which is new to the field. We found substantial inputs arising in the CA-regions of the hippocampus, which is a largely unexplored projection only documented in a few other studies (Wang et al., 2022; Zingg et al., 2018). Interestingly, CA1 inputs were not seen when rabies tracing was used with the Erg2-transgenic mouse line (Atlan et al., 2018), which could indicate that only a sub-population of CL cells receive inputs from these regions.

Although the true complexity of the CC might be best described by gradients rather than by defining clear borders (Atlan et al., 2017; Marriott et al., 2021; Olson & Graybiel, 1980) simple delineations still hold a practical value. We chose to provide a robust delineation system, based on a few easy-to-use chemical markers that can serve the communication and comparison of data between labs. Ultimately, our descriptive strategies should be determined by the anatomical precision needed to support particular experimental claims. In some cases, it will suffice to retrogradely label the CLRSC-pocket, whereas other experiments will call for multiple fiber- or cyto-architectural markers to be expressed. We showed that specific gene expression patterns aligned to anatomical features of both fiber- and cyto-architectural markers, together can be used as a common referencing system to anchor data in future experiments on the functional organization of the CC.

## Acknowledgements and funding

We would like to thank Qiangwei Zhang and Christina Schrick for assisting in the acquisition and processing of parts of the histological data. This project has received funding from the FriPro ToppForsk grant Enhanced Transgenics (90096000) of The Research Council of Norway to C.G.K., and supported by the Research Council of Norway through the Centre of Excellence scheme, and the National Infrastructure scheme (Centre for Neural Computation grant # 223262; NORBRAIN1 grant # 197467), and the Kavli Foundation to C.G.K. and M.P.W., the Wellcome Trust to A.M.P., the European Research Council (ERC) under the European Union’s Horizon 2020 research and innovation programme (grant agreement No 852765) to A.M.P., Natural Sciences and Engineering Research Council of Canada (NSERC) to D.K.O., and Clarendon Fund graduate scholar-ships to D.K.O. and A.M.S.

