## Supplementary tables 1 and 2 for "A multifaceted architectural framework of the mouse claustrum complex"

### Extended data

Table 7-1: Rabies tracing of monosynaptic (extrinsic) inputs to the claustrum complex

| Coarse categorizations | | Region | Input cells per region in animals Rb1-Rb13 | | | | | | | | | | | | |
|---|---|---|---|---|---|---|---|---|---|---|---|---|---|---|---|
| | | | Rb1 | Rb2 | Rb3 | Rb4 | Rb5 | Rb6 | Rb7 | Rb8 | Rb9 | Rb10 | Rb11 | Rb12 | Rb13 |
| Cortical | Prefrontal cortex | rIns | 38 | 20 | 7 | 89 | 15 | 22 | 7 | 4 | 10 | 10 | 10 | 19 | 8 |
| | | LO | 9 | 4 | 7 | 47 | 1 | 45 | 1 | 2 | 6 | 5 | 7 | 9 | 5 |
| | | VLO | 9 | 4 | 5 | 63 | 8 | 28 | 5 | 1 | 6 | 9 | 3 | 10 | 3 |
| | | VO | 2 | 2 | 3 | 17 | 0 | 17 | 2 | 0 | 4 | 7 | 0 | 1 | 0 |
| | | MO | 1 | 0 | 1 | 43 | 4 | 14 | 4 | 1 | 1 | 6 | 2 | 6 | 1 |
| | | PL | 3 | 2 | 4 | 66 | 2 | 31 | 5 | 1 | 4 | 6 | 5 | 9 | 4 |
| | | IL | 3 | 1 | 2 | 31 | 4 | 10 | 2 | 1 | 4 | 1 | 1 | 4 | 1 |
| | | D.Ped | 0 | 2 | 0 | 1 | 1 | 2 | 0 | 0 | 1 | 1 | 2 | 0 | 0 |
| | Cingulate cortex | dACC | 8 | 1 | 1 | 51 | 4 | 27 | 5 | 0 | 7 | 10 | 1 | 8 | 3 |
| | | vACC | 10 | 4 | 4 | 63 | 4 | 34 | 8 | 1 | 12 | 15 | 11 | 15 | 4 |
| | | RSC | 1 | 0 | 1 | 18 | 1 | 6 | 2 | 0 | 3 | 1 | 5 | 8 | 0 |
| | Parahippocampal region | LEC | 9 | 9 | 17 | 136 | 87 | 83 | 19 | 12 | 13 | 4 | 14 | 17 | 16 |
| | | MEC | 2 | 1 | 1 | 37 | 3 | 15 | 0 | 0 | 1 | 14 | 0 | 3 | 1 |
| | | PER | 3 | 1 | 10 | 34 | 2 | 11 | 23 | 9 | 0 | 2 | 3 | 2 | 8 |
| | | POR | 2 | 0 | 0 | 4 | 1 | 3 | 3 | 0 | 0 | 1 | 2 | 1 | 0 |
| | | PrS | 0 | 0 | 0 | 1 | 0 | 0 | 1 | 0 | 1 | 0 | 0 | 0 | 0 |
| | | PaS | 0 | 0 | 1 | 1 | 0 | 3 | 2 | 0 | 0 | 4 | 1 | 0 | 0 |
| | Hippocampal formation | CA1 | 11 | 1 | 5 | 36 | 12 | 40 | 3 | 2 | 5 | 3 | 6 | 16 | 7 |
| | | CA2 | 0 | 0 | 1 | 6 | 1 | 7 | 1 | 0 | 1 | 3 | 1 | 5 | 2 |
| | | CA3 | 0 | 0 | 0 | 43 | 0 | 0 | 0 | 0 | 0 | 0 | 0 | 1 | 0 |
| | | DG | 0 | 0 | 0 | 14 | 0 | 0 | 0 | 0 | 0 | 1 | 0 | 0 | 0 |
| | | Sub | 4 | 1 | 6 | 19 | 29 | 38 | 5 | 14 | 3 | 7 | 4 | 30 | 5 |
| | Sensorimotor cortex | SSc | 3 | 1 | 2 | 20 | 1 | 9 | 4 | 0 | 2 | 2 | 1 | 3 | 3 |
| | | Aud c | 2 | 0 | 1 | 7 | 0 | 4 | 1 | 1 | 1 | 1 | 0 | 1 | 0 |
| | | Vis c | 0 | 0 | 0 | 5 | 1 | 8 | 2 | 0 | 0 | 1 | 3 | 1 | 0 |
| | | M1 | 0 | 2 | 0 | 27 | 1 | 4 | 0 | 0 | 4 | 0 | 0 | 4 | 1 |
| | | rM2 | 0 | 1 | 1 | 19 | 2 | 3 | 0 | 0 | 0 | 1 | 1 | 2 | 4 |
| | | cM2 | 3 | 0 | 0 | 24 | 3 | 10 | 3 | 1 | 3 | 3 | 1 | 6 | 0 |
| | Other cortices | cIns | 4 | 7 | 12 | 14 | 11 | 12 | 20 | 4 | 10 | 1 | 2 | 6 | 4 |
| | | TeA | 1 | 0 | 1 | 10 | 0 | 2 | 2 | 0 | 1 | 3 | 0 | 4 | 0 |
| | | Par. c | 0 | 0 | 0 | 14 | 0 | 4 | 1 | 0 | 2 | 2 | 0 | 1 | 0 |
| Subcortical | Basal ganglia | CPu | 0 | 1 | 1 | 4 | 0 | 3 | 1 | 2 | 3 | 3 | 3 | 2 | 0 |
| | | GP | 3 | 0 | 0 | 14 | 3 | 5 | 0 | 1 | 0 | 2 | 2 | 0 | 0 |
| | | VP | 4 | 5 | 4 | 27 | 9 | 25 | 3 | 3 | 1 | 6 | 3 | 3 | 3 |
| | Basal forebrain | NAc | 0 | 1 | 0 | 3 | 0 | 3 | 0 | 0 | 0 | 2 | 1 | 4 | 0 |
| | | NDB | 10 | 9 | 22 | 77 | 23 | 39 | 1 | 8 | 1 | 14 | 15 | 17 | 3 |
| | | SIB | 2 | 0 | 2 | 5 | 0 | 11 | 0 | 0 | 0 | 0 | 0 | 0 | 0 |
| | Thalamus | Th MNG | 2 | 1 | 4 | 60 | 4 | 14 | 1 | 3 | 3 | 2 | 2 | 8 | 1 |
| | | Th ANG | 0 | 2 | 1 | 21 | 4 | 9 | 1 | 0 | 5 | 5 | 3 | 8 | 3 |
| | | Th LNG | 1 | 0 | 1 | 5 | 3 | 0 | 0 | 1 | 0 | 0 | 0 | 0 | 0 |
| | | Th VNG | 0 | 1 | 0 | 13 | 0 | 5 | 0 | 0 | 3 | 1 | 0 | 2 | 4 |
| | | Th IL | 1 | 2 | 0 | 19 | 2 | 11 | 0 | 0 | 2 | 3 | 0 | 1 | 2 |
| | | Th MD | 3 | 2 | 0 | 14 | 5 | 8 | 2 | 2 | 3 | 2 | 3 | 4 | 1 |
| | | MetaTh | 0 | 0 | 0 | 1 | 0 | 1 | 0 | 1 | 1 | 0 | 0 | 3 | 0 |
| | | Hb | 0 | 0 | 0 | 0 | 0 | 0 | 0 | 0 | 1 | 1 | 0 | 1 | 0 |
| | Hypothalamus | LH | 2 | 1 | 5 | 39 | 1 | 20 | 4 | 1 | 4 | 7 | 2 | 8 | 0 |
| | | MH | 1 | 0 | 0 | 8 | 0 | 5 | 1 | 0 | 1 | 0 | 2 | 3 | 0 |
| | | AH | 1 | 0 | 0 | 1 | 0 | 0 | 0 | 0 | 0 | 0 | 0 | 0 | 0 |
| | | PH | 1 | 0 | 0 | 5 | 0 | 2 | 1 | 0 | 0 | 1 | 0 | 1 | 0 |

| | | | | | | | | | | | | | | | |
|---|---|---|---|---|---|---|---|---|---|---|---|---|---|---|---|
| | Other subcortical areas | Sept | 1 | 0 | 0 | 1 | 1 | 10 | 0 | 0 | 1 | 0 | 1 | 2 | 2 |
| | | BNST | 0 | 0 | 0 | 1 | 0 | 3 | 0 | 0 | 0 | 0 | 0 | 0 | 0 |
| | | Preoptic | 0 | 0 | 0 | 5 | 2 | 2 | 0 | 0 | 0 | 1 | 0 | 0 | 0 |
| | | STh | 0 | 0 | 0 | 2 | 1 | 6 | 1 | 0 | 0 | 3 | 0 | 0 | 0 |
| | | ZI | 0 | 0 | 0 | 1 | 0 | 3 | 1 | 1 | 1 | 1 | 0 | 0 | 0 |
| | | Mam. n | 1 | 1 | 0 | 6 | 0 | 2 | 0 | 1 | 0 | 0 | 0 | 3 | 0 |
| Olfactory | Piriform area | rPir | 51 | 51 | 49 | 143 | 35 | 148 | 5 | 20 | 43 | 45 | 52 | 33 | 0 |
| | | cPir | 19 | 33 | 36 | 167 | 68 | 133 | 9 | 18 | 38 | 36 | 35 | 13 | 3 |
| | | Deep Pir | 5 | 4 | 5 | 23 | 1 | 11 | 2 | 3 | 3 | 2 | 3 | 2 | 0 |
| | | IEn | 4 | 11 | 5 | 22 | 5 | 15 | 2 | 0 | 3 | 10 | 9 | 3 | 0 |
| | | VEn | 0 | 15 | 1 | 7 | 0 | 9 | 1 | 5 | 3 | 6 | 2 | 2 | 0 |
| | Anterior olfactory area | AOL | 0 | 0 | 4 | 15 | 1 | 7 | 0 | 3 | 2 | 3 | 3 | 0 | 0 |
| | | AOM | 8 | 24 | 40 | 266 | 10 | 59 | 7 | 18 | 11 | 19 | 23 | 34 | 8 |
| | | AOV/P | 2 | 7 | 7 | 29 | 2 | 22 | 0 | 4 | 3 | 3 | 6 | 5 | 1 |
| | | AOD | 0 | 2 | 1 | 19 | 2 | 2 | 0 | 1 | 2 | 1 | 2 | 2 | 0 |
| | Other olfactory areas | VTT | 1 | 2 | 6 | 19 | 2 | 23 | 1 | 5 | 3 | 4 | 5 | 4 | 0 |
| | | DTT | 13 | 2 | 7 | 24 | 9 | 17 | 5 | 4 | 2 | 6 | 8 | 4 | 0 |
| | | Tu | 0 | 0 | 0 | 1 | 1 | 7 | 0 | 0 | 1 | 1 | 0 | 0 | 0 |
| | | LOT | 2 | 2 | 4 | 10 | 1 | 11 | 1 | 5 | 1 | 4 | 3 | 3 | 0 |
| | | OB | 0 | 0 | 0 | 1 | 0 | 0 | 0 | 0 | 0 | 1 | 0 | 0 | 0 |
| Amygdala | Cortical amygdala | AA | 1 | 4 | 6 | 25 | 3 | 24 | 2 | 6 | 4 | 12 | 3 | 3 | 0 |
| | | CxA | 4 | 2 | 15 | 33 | 10 | 26 | 5 | 12 | 8 | 4 | 10 | 3 | 1 |
| | | IPAC | 2 | 0 | 0 | 9 | 1 | 5 | 1 | 5 | 0 | 2 | 4 | 1 | 0 |
| | | ACo | 1 | 0 | 0 | 9 | 2 | 7 | 0 | 2 | 1 | 0 | 0 | 0 | 0 |
| | | PCo | 1 | 5 | 9 | 24 | 7 | 23 | 3 | 9 | 2 | 2 | 3 | 8 | 1 |
| | | APir | 2 | 3 | 14 | 30 | 12 | 15 | 2 | 9 | 2 | 3 | 2 | 12 | 1 |
| | Basolateral complex | BLA | 7 | 6 | 3 | 133 | 11 | 57 | 4 | 6 | 2 | 16 | 4 | 21 | 3 |
| | | BLP | 5 | 2 | 2 | 22 | 6 | 9 | 3 | 2 | 0 | 1 | 0 | 2 | 1 |
| | | BLV | 0 | 0 | 0 | 0 | 0 | 2 | 0 | 0 | 0 | 0 | 1 | 0 | 0 |
| | | LA | 1 | 1 | 1 | 11 | 1 | 2 | 1 | 2 | 0 | 1 | 0 | 3 | 0 |
| | | BMA | 3 | 0 | 4 | 27 | 2 | 9 | 2 | 0 | 1 | 1 | 1 | 2 | 0 |
| | Centromedial Amyg-dala | CeA | 0 | 0 | 1 | 0 | 0 | 1 | 0 | 2 | 1 | 0 | 0 | 0 | 0 |
| | | Med A | 2 | 0 | 5 | 40 | 3 | 29 | 5 | 4 | 4 | 4 | 1 | 7 | 0 |
| Brainstem | Brainstem | PAG | 0 | 0 | 0 | 3 | 0 | 5 | 3 | 1 | 0 | 0 | 1 | 0 | 0 |
| | | SN | 0 | 0 | 0 | 1 | 0 | 0 | 1 | 0 | 0 | 0 | 1 | 0 | 0 |
| | | VTA | 0 | 0 | 0 | 3 | 0 | 1 | 0 | 0 | 0 | 1 | 1 | 0 | 0 |
| | | Rn | 3 | 3 | 3 | 30 | 3 | 25 | 3 | 2 | 4 | 1 | 8 | 5 | 0 |
| | | Rt | 0 | 0 | 0 | 7 | 0 | 1 | 1 | 0 | 0 | 1 | 0 | 0 | 0 |
| | | Pn | 0 | 0 | 1 | 6 | 0 | 3 | 0 | 0 | 0 | 0 | 1 | 0 | 0 |
| | | Other BS | 0 | 0 | 2 | 7 | 0 | 5 | 0 | 2 | 1 | 0 | 1 | 0 | 0 |
| | | Total | 298 | 269 | 364 | 2458 | 454 | 1392 | 217 | 228 | 280 | 368 | 316 | 434 | 118 |

### Table 7-2: Contralateral inputs in rabies tracing dataset

| Coarse categorizations | | Region | Contralateral input cells per region in animals Rb1-Rb13 | | | | | | | | | | | | |
|---|---|---|---|---|---|---|---|---|---|---|---|---|---|---|---|
| | | | Rb1 | Rb2 | Rb3 | Rb4 | Rb5 | Rb6 | Rb7 | Rb8 | Rb9 | Rb10 | Rb11 | Rb12 | Rb13 |
| Cortical | Prefrontal cortex | rIns | 0 | 0 | 0 | 1 | 0 | 1 | 0 | 0 | 0 | 0 | 0 | 2 | 0 |
| | | LO | 0 | 0 | 0 | 4 | 1 | 4 | 0 | 0 | 0 | 0 | 1 | 0 | 0 |
| | | VLO | 0 | 1 | 1 | 15 | 2 | 4 | 1 | 0 | 0 | 0 | 1 | 3 | 0 |
| | | VO | 1 | 0 | 0 | 2 | 0 | 1 | 1 | 0 | 1 | 3 | 0 | 0 | 0 |
| | | MO | 1 | 0 | 1 | 11 | 2 | 2 | 1 | 1 | 0 | 0 | 0 | 4 | 1 |
| | | PL | 2 | 1 | 2 | 36 | 1 | 16 | 3 | 1 | 1 | 5 | 3 | 4 | 2 |
| | | IL | 2 | 0 | 2 | 9 | 4 | 2 | 1 | 0 | 1 | 0 | 1 | 0 | 0 |
| | | DPed | 0 | 0 | 0 | 1 | 0 | 0 | 0 | 0 | 0 | 0 | 0 | 0 | 0 |
| | Cingulate cortex | dACC | 4 | 0 | 0 | 33 | 1 | 11 | 4 | 0 | 5 | 5 | 1 | 6 | 1 |
| | | vACC | 3 | 2 | 2 | 33 | 2 | 20 | 5 | 0 | 7 | 7 | 7 | 8 | 1 |
| | | RSp | 0 | 0 | 0 | 0 | 0 | 0 | 0 | 0 | 0 | 0 | 1 | 1 | 0 |
| | Parahippocampal region | LEC | 0 | 0 | 1 | 2 | 2 | 4 | 0 | 0 | 1 | 0 | 0 | 0 | 0 |
| | | MEC* | 0 | 0 | 1 | 12 | 1 | 0 | 0 | 0 | 0 | 0 | 0 | 0 | 0 |
| | | PER | 0 | 0 | 1 | 1 | 0 | 2 | 0 | 0 | 0 | 1 | 0 | 0 | 0 |
| | | POR* | 0 | 0 | 0 | 1 | 0 | 0 | 0 | 0 | 0 | 0 | 0 | 0 | 0 |
| | Hippocampal formation | CA1 | 0 | 0 | 0 | 0 | 0 | 0 | 1 | 0 | 0 | 0 | 0 | 0 | 0 |
| | | CA2 | 0 | 0 | 0 | 0 | 0 | 0 | 0 | 0 | 1 | 0 | 0 | 0 | 0 |
| | | CA3 | 0 | 0 | 0 | 12 | 0 | 0 | 0 | 0 | 0 | 0 | 0 | 0 | 0 |
| | | DG | 0 | 0 | 0 | 0 | 0 | 0 | 0 | 0 | 0 | 1 | 0 | 0 | 0 |
| | | Sub | 0 | 0 | 0 | 0 | 1 | 0 | 0 | 1 | 0 | 0 | 0 | 0 | 0 |
| | Sensorimotor cortex | SSc | 0 | 0 | 1 | 6 | 0 | 3 | 2 | 0 | 0 | 0 | 0 | 1 | 0 |
| | | Aud c | 0 | 0 | 0 | 1 | 0 | 0 | 0 | 0 | 0 | 1 | 0 | 0 | 0 |
| | | Vis c | 0 | 0 | 0 | 0 | 0 | 1 | 0 | 0 | 0 | 1 | 0 | 0 | 0 |
| | | M1 | 0 | 1 | 0 | 14 | 0 | 1 | 0 | 0 | 2 | 0 | 0 | 1 | 1 |
| | | rM2 | 0 | 1 | 0 | 15 | 0 | 2 | 0 | 0 | 0 | 0 | 0 | 2 | 0 |
| | | cM2 | 2 | 0 | 0 | 12 | 1 | 5 | 2 | 1 | 2 | 2 | 1 | 3 | 0 |
| | Other cortices | TeA | 0 | 0 | 1 | 0 | 0 | 0 | 0 | 0 | 0 | 0 | 0 | 0 | 0 |
| | | Par c | 0 | 0 | 0 | 1 | 0 | 0 | 0 | 0 | 0 | 0 | 0 | 0 | 0 |
| Subcortical | Basal forebrain | NAc | 0 | 0 | 0 | 3 | 0 | 0 | 0 | 0 | 0 | 1 | 1 | 0 | 0 |
| | | NDB | 1 | 1 | 0 | 1 | 0 | 2 | 0 | 0 | 0 | 1 | 0 | 1 | 0 |
| | Thalamus | Th ANG | 0 | 0 | 0 | 0 | 0 | 0 | 0 | 0 | 0 | 0 | 0 | 1 | 0 |
| | | Th IL | 0 | 0 | 0 | 1 | 0 | 0 | 0 | 0 | 0 | 0 | 0 | 0 | 0 |
| | | Th MD | 0 | 0 | 0 | 1 | 0 | 0 | 0 | 1 | 0 | 0 | 0 | 0 | 0 |
| | | MetaTh | 0 | 0 | 0 | 0 | 0 | 0 | 0 | 1 | 0 | 0 | 0 | 0 | 0 |
| | Hypothalamus | LH | 0 | 0 | 0 | 4 | 0 | 2 | 1 | 0 | 1 | 1 | 0 | 0 | 0 |
| | | MH | 0 | 0 | 0 | 0 | 0 | 0 | 0 | 0 | 0 | 0 | 0 | 1 | 0 |
| | Other subcortical areas | ZI | 0 | 0 | 0 | 0 | 0 | 1 | 0 | 0 | 0 | 0 | 0 | 0 | 0 |
| Olfactory | Piriform area | rPir | 0 | 0 | 2 | 6 | 0 | 7 | 0 | 0 | 0 | 1 | 1 | 1 | 0 |
| | | cPir | 0 | 0 | 2 | 2 | 0 | 0 | 0 | 0 | 0 | 0 | 2 | 0 | 1 |
| | Anterior olfactory area | AOL | 0 | 0 | 0 | 2 | 0 | 5 | 0 | 0 | 0 | 0 | 1 | 0 | 0 |
| | | AOM | 1 | 5 | 1 | 33 | 1 | 10 | 0 | 2 | 6 | 5 | 6 | 3 | 3 |
| | | AOV/P | 1 | 2 | 1 | 12 | 1 | 7 | 0 | 0 | 2 | 0 | 3 | 0 | 0 |
| | Other olfactory areas | DTT | 0 | 0 | 0 | 0 | 0 | 0 | 0 | 0 | 0 | 1 | 1 | 0 | 0 |
| | | LOT | 1 | 0 | 0 | 2 | 0 | 2 | 0 | 0 | 0 | 2 | 1 | 0 | 0 |
| Amygdala | Cortical amygdala | AA | 1 | 1 | 0 | 2 | 0 | 0 | 0 | 1 | 0 | 0 | 0 | 0 | 0 |
| | | PCo | 0 | 0 | 1 | 0 | 0 | 0 | 0 | 0 | 0 | 0 | 0 | 0 | 0 |
| | Basolateral complex | BLA | 2 | 0 | 0 | 29 | 2 | 7 | 1 | 2 | 2 | 4 | 0 | 1 | 0 |
| | | BLP | 1 | 1 | 0 | 2 | 0 | 0 | 0 | 0 | 0 | 0 | 0 | 0 | 0 |
| | | BLV | 0 | 0 | 0 | 0 | 0 | 1 | 0 | 0 | 0 | 0 | 0 | 0 | 0 |
| | | LA | 0 | 1 | 0 | 0 | 0 | 0 | 0 | 0 | 0 | 0 | 0 | 0 | 0 |
| | | BMA | 0 | 0 | 3 | 2 | 0 | 1 | 0 | 0 | 0 | 0 | 0 | 1 | 0 |
| BS | Brainstem | Rt | 0 | 0 | 0 | 2 | 0 | 0 | 0 | 0 | 0 | 1 | 0 | 0 | 0 |
| | | Pn | 0 | 0 | 0 | 2 | 0 | 1 | 0 | 0 | 0 | 0 | 1 | 0 | 0 |
| | | Sum | 23 | 17 | 23 | 328 | 22 | 125 | 23 | 11 | 32 | 43 | 33 | 44 | 10 |

* Inputs from MEC and POR could not be unequivocally classified as ipsi- or contralateral since the hemispheres were detached from the rest of the tissue in the caudal-most parts of the brain.
